# Simple Framework for Constructing Functional Spiking Recurrent Neural Networks

**DOI:** 10.1101/579706

**Authors:** Robert Kim, Yinghao Li, Terrence J. Sejnowski

## Abstract

Cortical microcircuits exhibit complex recurrent architectures that possess dynamically rich properties. The neurons that make up these microcircuits communicate mainly via discrete spikes, and it is not clear how spikes give rise to dynamics that can be used to perform computationally challenging tasks. In contrast, continuous models of rate-coding neurons can be trained to perform complex tasks. Here, we present a simple framework to construct biologically realistic spiking recurrent neural networks (RNNs) capable of learning a wide range of tasks. Our framework involves training a continuous-variable rate RNN with important biophysical constraints and transferring the learned dynamics and constraints to a spiking RNN in a one-to-one manner. The proposed framework introduces only one additional parameter to establish the equivalence between rate and spiking RNN models. We also study other model parameters related to the rate and spiking networks to optimize the one-to-one mapping. By establishing a close relationship between rate and spiking models, we demonstrate that spiking RNNs could be constructed to achieve similar performance as their counterpart continuous rate networks.

## Introduction

Dense recurrent connections common in cortical circuits suggest their important role in computational processes [1–3]. Network models based on recurrent neural networks (RNNs) of continuous-variable rate units have been extensively studied to characterize network dynamics underlying neural computations [4–9]. Methods commonly used to train rate networks to perform cognitive tasks can be largely classified into three categories: recursive least squares (RLS)-based, gradient-based, and reward-based algorithms. The First-Order Reduced and Controlled Error (FORCE) algorithm, which utilizes RLS, has been widely used to train RNNs to produce complex output signals [5] and to reproduce experimental results [6, 10, 11]. Gradient descent-based methods, including Hessian-free methods, have been also successfully applied to train rate networks in a supervised manner and to replicate the computational dynamics observed in networks from behaving animals [7, 12, 13]. Unlike the previous two categories (i.e. RLS-based and gradient-based algorithms), reward-based learning methods are more biologically plausible and have been shown to be as effective in training rate RNNs as the supervised learning methods [14–17]. Even though these models have been vital in uncovering previously unknown computational mechanisms, continuous rate networks do not incorporate basic biophysical constraints such as the spiking nature of biological neurons.

Training spiking network models where units communicate with one another via discrete spikes is more difficult than training continuous rate networks. The non-differentiable nature of spike signals prevents the use of gradient descent-based methods to train spiking networks directly, although several differentiable models have been proposed [18, 19]. Due to this challenge, FORCE-based learning algorithms have been most commonly used to train spiking recurrent networks. While recent advances have successfully modified and applied FORCE training to construct functional spike RNNs [8, 20–23], FORCE training is computationally inefficient and unstable when connectivity constraints, including separate populations for excitatory and inhibitory populations (Dale’s principle) and sparse connectivity patterns, are imposed [21].

Due to these limitations, computational capabilities of spiking networks that abide by biological constraints have been challenging to explore. For instance, it is not clear if spiking RNNs operating in a purely rate-coding regime can perform tasks as complex as the ones rate RNN models are trained to perform. If such spiking networks can be constructed, then it would be important to characterize how much spiking-related noise not present in rate networks affects the performance of the networks. Establishing the relationship between these two types of RNN models could also serve as a good starting point for designing power-efficient spiking networks that can incorporate both rate and temporal coding.

To address the above questions, we present a computational framework for directly mapping rate RNNs with basic biophysical constraints to leaky integrate-and-fire (LIF) spiking RNNs without significantly compromising task performance. Our method introduces only one additional parameter to place the spiking RNNs in the same dynamic regime as their counterpart rate RNNs, and takes advantage of the previously established methods to efficiently optimize network parameters while adhering to biophysical restrictions. These previously established methods include training a continuous-variable rate RNN using a gradient descent-based method [24–27] and connectivity weight matrix parametrization method to impose Dale’s principle [13]. The gradient descent learning algorithm allowed us to easily optimize many parameters including the connectivity weights of the network and the synaptic decay time constant for each unit. The weight parametrization method proposed by Song et al. was utilized to enforce Dale’s principles and additional connectivity patterns without significantly affecting computational efficiency and network stability [13].

Combining these two existing methods with correct parameter values enabled us to directly map rate RNNs trained with backpropagation to LIF RNNs in a one-to-one manner. The parameters critical for mapping to succeed included the network size, the nonlinear activation function employed for training rate RNNs, and a constant factor for scaling down the connectivity weights of the trained rate RNNs. Here, we investigated these parameters along with other LIF parameters and identified the range of values required for the mapping to be effective. We demonstrate that when these parameters are set to their optimal values, the LIF models constructed from our framework can perform the same tasks the rate models are trained to perform equally well.

## Results

Here we provide a brief overview of the two types of recurrent neural networks (RNNs) that we employed throughout this study (more details in Methods): continuous-variable firing rate RNNs and spiking RNNs. The continuous-variable rate network model consisted of *N* rate units whose firing rates were estimated via a nonlinear input-output transfer function [4, 5]. The model was governed by the following set of equations:

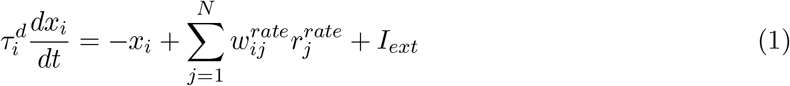

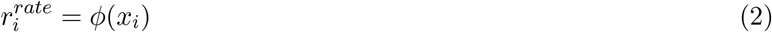

where 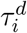 is the synaptic decay time constant for unit *i*, *x*_*i*_ is the synaptic current variable for unit *i*, 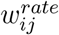 is the synaptic strength from unit *j* to unit *i*, and *I*_*ext*_ is the external current input to unit *i*. The firing rate of unit 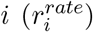 is given by applying a nonlinear transfer function (*φ*(⋅)) to the synaptic current variable. Since the firing rates in spiking networks cannot be negative, we chose the activation function for our rate networks to be a non-negative saturating function (standard sigmoid function) and parametrized the connectivity matrix 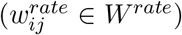 to enforce Dale’s principle and additional connectivity constraints (see Methods).

The second RNN model that we considered was a network composed of *N* spiking units. Throughout this study, we focused on networks of leaky integrate-and-fire (LIF) units whose membrane voltage dynamics were given by:

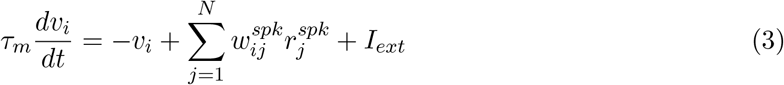

where *τ*_*m*_ is the membrane time constant (set to 10 ms throughout this study), *v*_*i*_ is the membrane voltage of unit *i*, 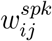 is the synaptic strength from unit *j* to unit *i*, 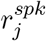 represents the synaptic filtering of the spike train of unit *j*, and *I*_*ext*_ is the external current source. The discrete nature of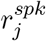 (see Methods) has posed a major challenge for directly training spiking networks using gradient-based supervised learning. Even though the main results presented here are based on LIF networks, our method can be generalized to quadratic integrate-and-fire (QIF) networks with only few minor changes to the model parameters (SI Appendix, Table S1).

Continuous rate network training was implemented using the open-source software library TensorFlow in Python, while LIF/QIF network simulations along with the rest of the analyses were performed in MATLAB.

### Training Continuous Rate Networks

Throughout this study, we used a gradient-descent supervised method, known as Backpropagation Through Time (BPTT), to train rate RNNs to produce target signals associated with a specific task [13, 24]. The method we employed is similar to the one used by previous studies ([13, 25, 27]; more details in Methods) with one major difference in synaptic decay time constants. Instead of assigning a single time constant to be shared by all the units in a network, our method tunes a synaptic constant for each unit using BPTT (see Methods). Although tuning of synaptic time constants may not be biologically plausible, this feature was included to model diverse intrinsic synaptic timescales observed in single cortical neurons [28–30].

We trained rate RNNs of various sizes on a simple task modeled after a Go-NoGo task to demonstrate our training method (Fig. 1). Each network was trained to produce a positive mean population activity approaching +1 after a brief input pulse (Fig. 1A). For a trial without an input pulse (i.e. NoGo trial), the networks were trained to maintain the output signal close to zero. The units in a rate RNN were sparsely connected via *W*^*rate*^ and received a task-specific input signal through weights (*W*_*in*_) drawn from a normal distribution with zero mean and unit variance. The network output (*o*^*rate*^) was then computed using a set of linear readout weights:

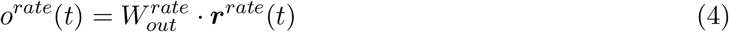

where 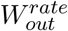 is the readout weights and ***r***^*rate*^(*t*) is the firing rate estimates from all the units in the network at time *t*. The recurrent weight matrix (*W*^*rate*^), the readout weights 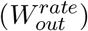, and the synaptic decay time constants (***τ***^***d***^) were optimized during training, while the input weight matrix (*W*_*in*_) stayed fixed (see Methods).

**Fig. 1.**
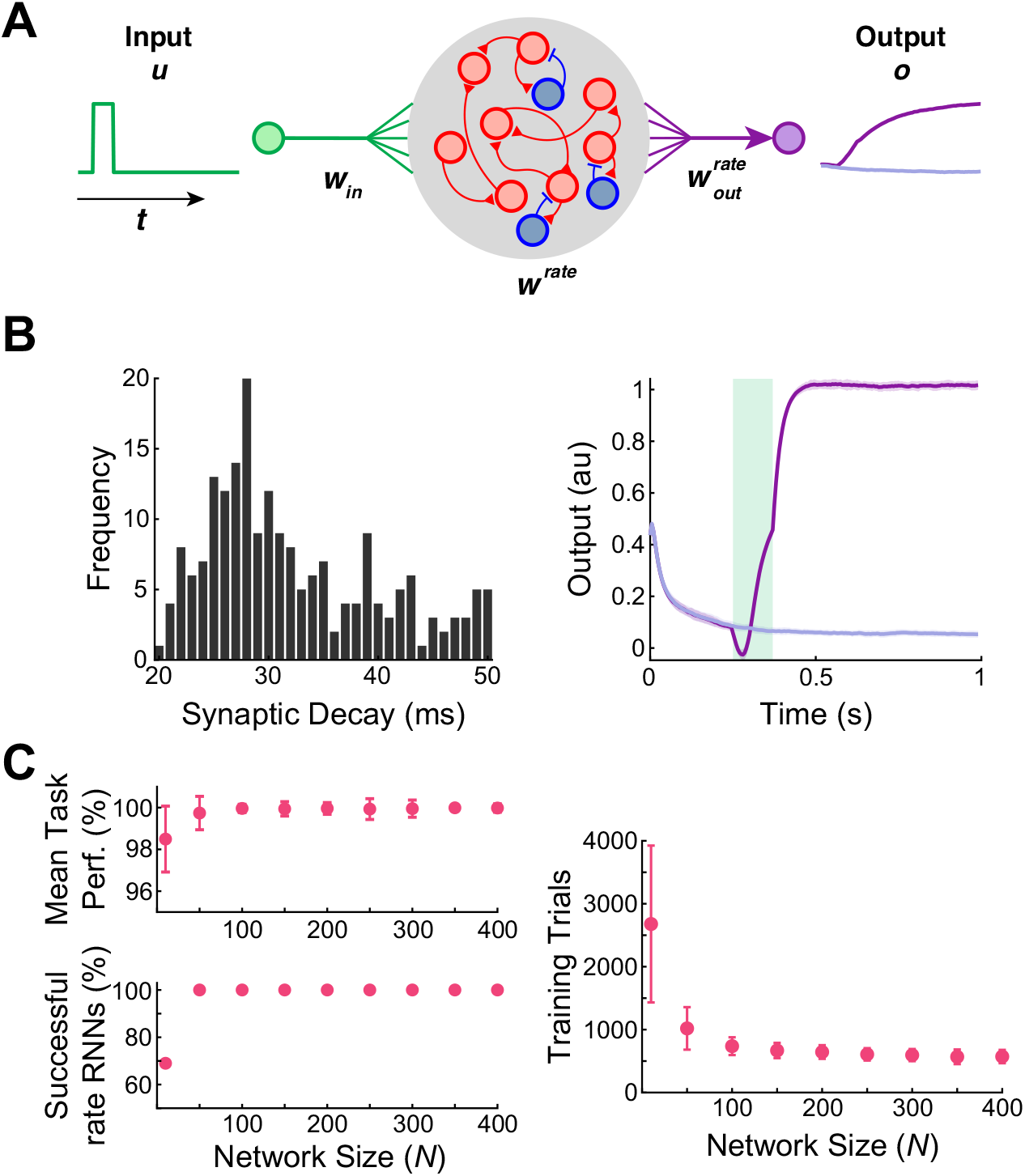
Rate RNNs trained to perform the Go-NoGo task. **A.** Schematic diagram illustrating a continuous rate RNN model trained to perform the Go-NoGo task. The rate RNN model contained excitatory (red circles) and inhibitory (blue circles) units. **B.** Distribution of the tuned synaptic decay time constants (Mean ± SD, 28.2 ± 9.4 ms; left) and the average trained rate RNN task performance (right) from an example rate RNN model. The mean ± SD output signals from 50 Go trials (dark purple) and from 50 NoGo trials (light purple) are shown. The green box represents the input stimulus given for the Go trials. The rate RNN contained 200 units (169 excitatory and 31 inhibitory units). **C.** Rate RNNs with different network sizes trained to perform the Go-NoGo task. For each network size, 100 RNNs with random initial conditions were trained. All the networks successfully trained performed the task almost perfectly (range 96–100%; left). As the network size increased, the number of training trials decreased (Mean ± SD shown; right).

The network size (*N*) was varied from 10 to 400 (9 different sizes), and 100 networks with random initializations were trained for each size. For all the networks, the minimum and the maximum synaptic decay time constants were fixed to 20 ms and 50 ms, respectively. As expected, the smallest rate RNNs (*N* = 10) took the longest to train, and only 69% of the rate networks with *N* = 10 were successfully trained (see SI Appendix for training termination criteria; Fig. 1C).

### One-to-One Mapping from Continuous Rate Networks to Spiking Networks

We developed a simple procedure that directly maps dynamics of a trained continuous rate RNN to a spiking RNN in a one-to-one manner.

In our framework, the three sets of the weight matrices (*W*_*in*_, *W*^*rate*^, and 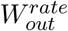) along with the tuned synaptic time constants (***τ***^***d***^) from a trained rate RNN are transferred to a network of LIF spiking units. The spiking RNN is initialized to have the same topology as the rate RNN. The input weight matrix and the synaptic time constants are simply transferred without any modification, but the recurrent connectivity and the readout weights need to be scaled by a constant factor (*λ*) in order to account for the difference in the firing rate scales between the rate model and the spiking model (see Methods; Fig. 2A). The effects of the scaling factor is clear in an example LIF RNN model constructed from a rate model trained to perform the Go-NoGo task (Fig. 2B). With an appropriate value for *λ*, the LIF network performed the task with the same accuracy as the rate network, and the LIF units fired at rates similar to the “rates” of the continuous network units (SI Appendix, Fig. S1). In addition, the LIF network reproduced the population dynamics of the rate RNN model as shown by the time evolution of the top three principal components extracted by the principal component analysis (SI Appendix, Fig. S2).

**Fig. 2.**
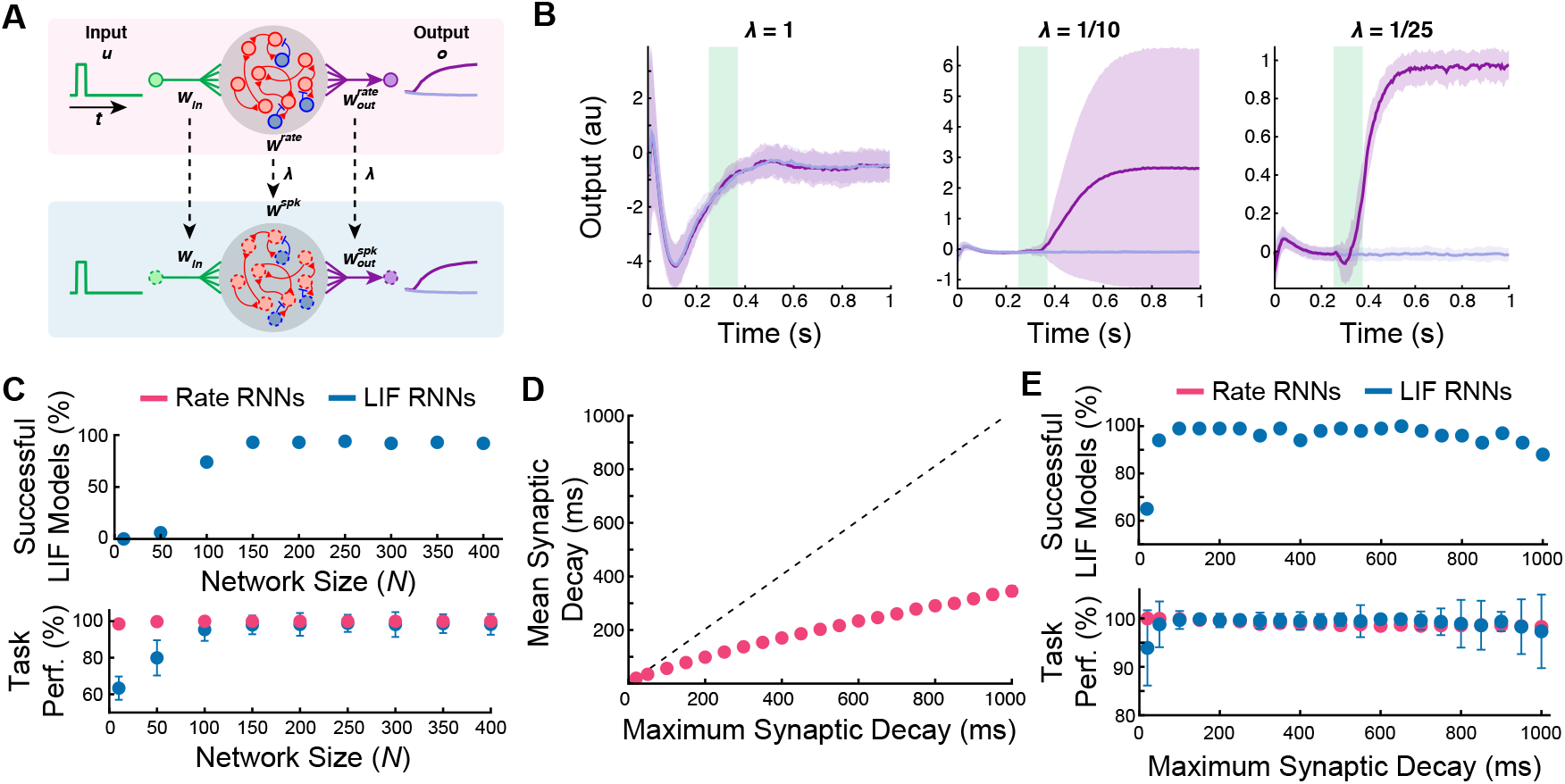
Mapping trained rate RNNs to LIF RNNs for the Go-NoGo task. **A.** Schematic diagram illustrating direct mapping from a continuous rate RNN model (top) to a spiking RNN model (bottom). The optimized synaptic decay time constants (***τ***^***d***^) along with the weight parameters (*W*_*in*_, *W*^*rate*^, and 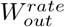) were transferred to a spiking network with LIF units (red and blue circles with a dashed outline). The connectivity and the readout weights were scaled by a constant factor, *λ*. **B.** LIF RNN performance on the Go-NoGo task without scaling (*λ* = 1; left), with insufficient scaling (middle), and with appropriate scaling (right). The network contained 200 units (169 excitatory and 31 inhibitory units). Mean ± SD over 50 Go and 50 NoGo trials. **C.** Successfully converted LIF networks and their average task performance on the Go-NoGo task with different network sizes. All the rate RNNs trained in Fig. 1 were converted to LIF RNNs. The network size was varied from *N* = 10 to 400. **D.** Average synaptic decay values for *N* = 250 across different maximum synaptic decay constants. **E.** Successfully converted LIF networks and their average task performance on the Go-NoGo task with fixed network size (*N* = 250) and different maximum synaptic decay constants. The maximum synaptic decay constants were varied from 20 ms to 1000 ms.

Using the procedure outlined above, we converted all the rate RNNs trained in the previous section to spiking RNNs. Only the rate RNNs that successfully performed the task (i.e. training termination criteria met within the first 6000 trials) were converted. Fig. 2C characterizes the proportion of the LIF networks that successfully performed the Go-NoGo task (≥ 95% accuracy; same threshold used to train the rate models; see SI Appendix) and the average task performance of the LIF models for each network size group. For each conversion, the scaling factor (*λ*) was determined via a grid search method (see Methods). The LIF RNNs constructed from the small rate networks (*N* = 10 and *N* = 50) did not perform the task reliably, but the LIF model became more robust as the network size increased, and the performance gap between the rate RNNs and the LIF RNNs was the smallest for *N* = 250 (Fig. 2C).

In order to investigate the effects of the synaptic decay time constants on the mapping robustness, we trained rate RNNs composed of 250 units (*N* = 250) with different maximum time constants 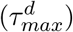. The minimum time constant 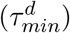 was fixed to 20 ms, while the maximum constant was varied from 20 ms to 1000 ms. For the first case (i.e.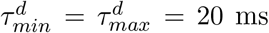), the synaptic decay time constants were not trained and fixed to 20 ms for all the units in a rate RNN. For each maximum constant value, 100 rate RNNs with different initial conditions were trained, and only successfully trained rate networks were converted to spiking RNNs. For each maximum synaptic decay condition, all 100 rate RNNs were successfully trained. As the maximum decay constant increased, the average tuned synaptic decay constants increased sub-linearly (Fig. 2D). For the shortest synaptic decay time constant considered (20 ms), the average task performance was the lowest at 93.91 ± 7.78%, and 65% of the converted LIF RNNs achieved at least 95% accuracy (Fig. 2E). The LIF models for the rest of the maximum synaptic decay conditions were robust. Although this might indicate that tuning of ***τ***^***d***^ is important for the conversion of rate RNNs to LIF RNNs, we further investigated the effects of the optimization of ***τ***^***d***^ in the last section (see Analysis of the Conversion Method).

Our framework also allows seamless integration of additional functional connectivity constraints. For example, a common cortical microcircuitry motif where somatostatin-expressing interneurons inhibit both pyramidal and parvalbumin-positive neurons can be easily implemented in our framework (see Methods and SI Appendix, Fig. S3). In addition, Dale’s principle is not required for our framework (SI Appendix, Fig. S4).

### LIF networks for context-dependent input integration

The Go-NoGo task considered in the previous section did not require complex cognitive computations. In this section, we consider a more complex task and probe whether spiking RNNs can be constructed from trained rate networks in a similar fashion. The task considered here is modeled after the context-dependent sensory integration task employed by Mante et al. [7]. Briefly, Mante et al. trained rhesus monkeys to integrate inputs from one sensory modality (dominant color or dominant motion of randomly moving dots) while ignoring inputs from the other modality [7]. A contextual cue was also given to instruct the monkeys which sensory modality they should attend to. The task required the monkeys to utilize flexible computations as the same modality can be either relevant or irrelevant depending on the contextual cue. Previous works have successfully trained continuous rate RNNs to perform a simplified version of the task and replicated the neural dynamics present in the experimental data [7, 13, 15]. Using our framework, we constructed the first spiking RNN model to our knowledge that can perform the task and capture the dynamics observed in the experimental data.

For the task paradigm, we adopted a similar design as the one used by the previous modeling studies [7, 13, 15]. A network of recurrently connected units received two streams of noisy input signals along with a constant-valued signal that encoded the contextual cue (Fig. 3A; see Methods). To simulate a noisy sensory input signal, a random Gaussian time-series signal with zero mean and unit variance was first generated. Each input signal was then shifted by a positive or negative constant (“offset”) to encode evidence toward the (+) or (-) choice, respectively. Therefore, the offset value determined how much evidence for the specific choice was represented in the noisy input signal. The network was trained to produce an output signal approaching +1 (or −1) if the cued input signal had a positive (or negative) mean. For example, if the cued input signal was generated using a positive offset value, then the network should produce an output that approaches +1 regardless of the mean of the irrelevant input signal.

**Fig. 3.**
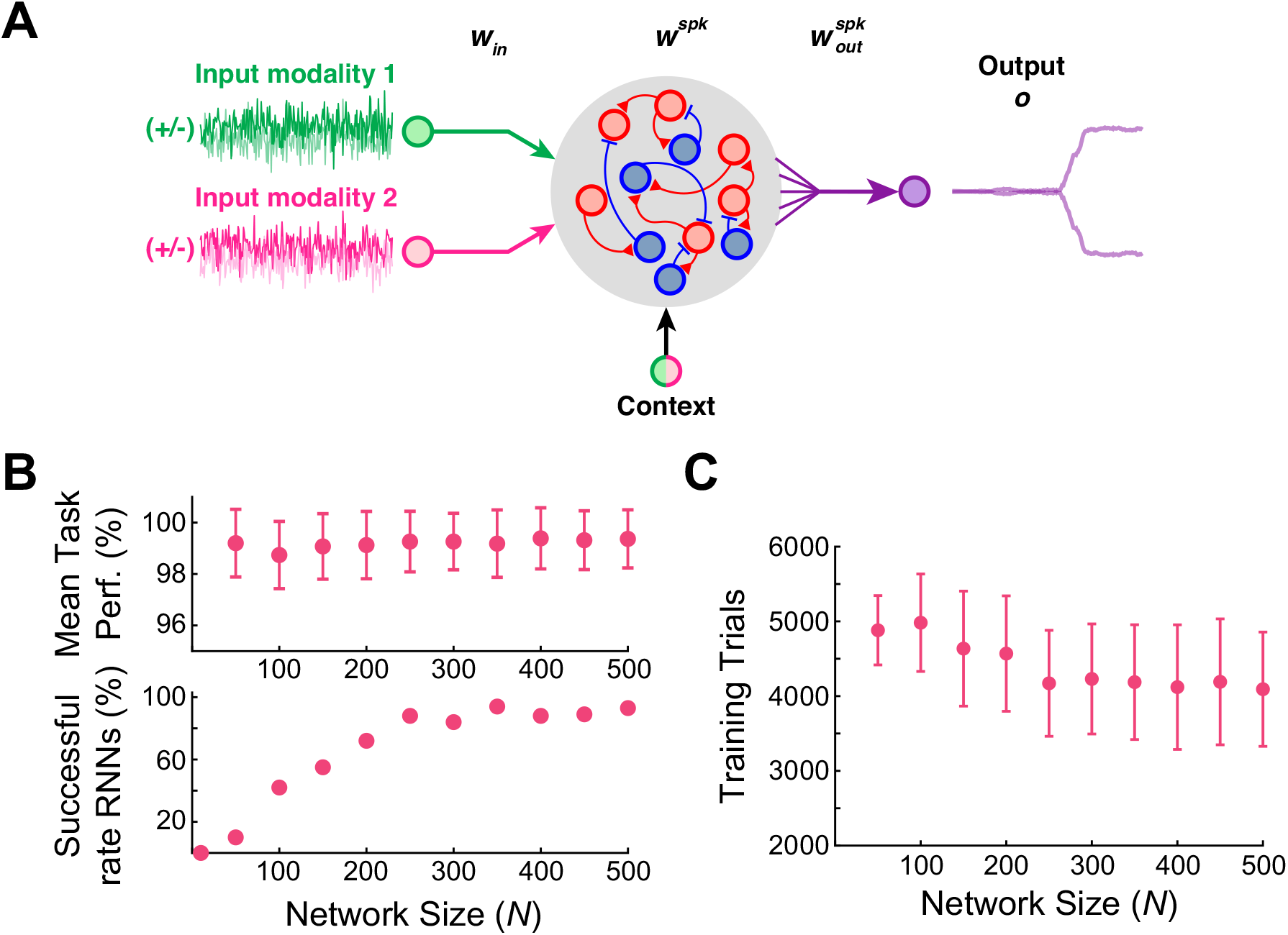
Rate RNNs trained to perform the contextual integration task. **A.** Diagram illustrating the task paradigm modeled after the context-dependent task used by Mante et al. [7]. Two streams of noisy input signals (green and magenta lines) along with a context signal were delivered to the LIF network. The network was trained to integrate and determine if the mean of the cued input signal (i.e. cued offset value) was positive (“+” choice) or negative (“-” choice). **B.** Rate RNNs with different network sizes trained to perform the contextual integration task. The network size was varied from *N* = 10 to 500. For each network size, 100 RNNs with random initial conditions were trained. The average task performance (top) and the proportion of the successful rate models (bottom) are shown. A model was successful if its mean task performance was ≥ 95%. **C.** Average number of training trials required for each network size. As the network size increased, the number of training trials decreased (Mean ± SD shown).

Rate networks with different sizes (*N* = 10, 50, …, 450, 500) were trained to perform the task. As this is a more complex task compared to the Go-NoGo task considered in the previous section, the number of units and trials required to train rate RNNs was larger than the models trained on the Go-NoGo task (Fig. 3B and 3C). The synaptic decay time constants were again limited to a range of 20 ms and 50 ms, and 100 rate RNNs with random initial conditions were trained for each network size. For the smallest network size (*N* = 10), the rate networks could not be trained to perform the task within the first 6000 trials (Fig. 3B).

Next, all the rate networks successfully trained for the task were transformed into LIF models. Example output responses along with the distribution of the tuned synaptic decay constants from a converted LIF model (*N* = 250, 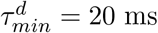, 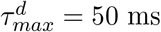) are shown in Fig. 4A and 4B. The task performance of the LIF model was 98% and comparable to the rate RNN used to construct the spiking model (Fig. 4C). In addition, the LIF network manifested population dynamics similar to the dynamics observed in the group of neurons recorded by Mante et al. [7] and rate RNN models investigated in previous studies [7, 13, 15]: individual LIF units displayed mixed representation of the four task variables (modality 1, modality 2, network choice, and context; see SI Appendix, Fig. S5A), and the network revealed the characteristic line attractor dynamics (SI Appendix, Fig. S5B).

**Fig. 4.**
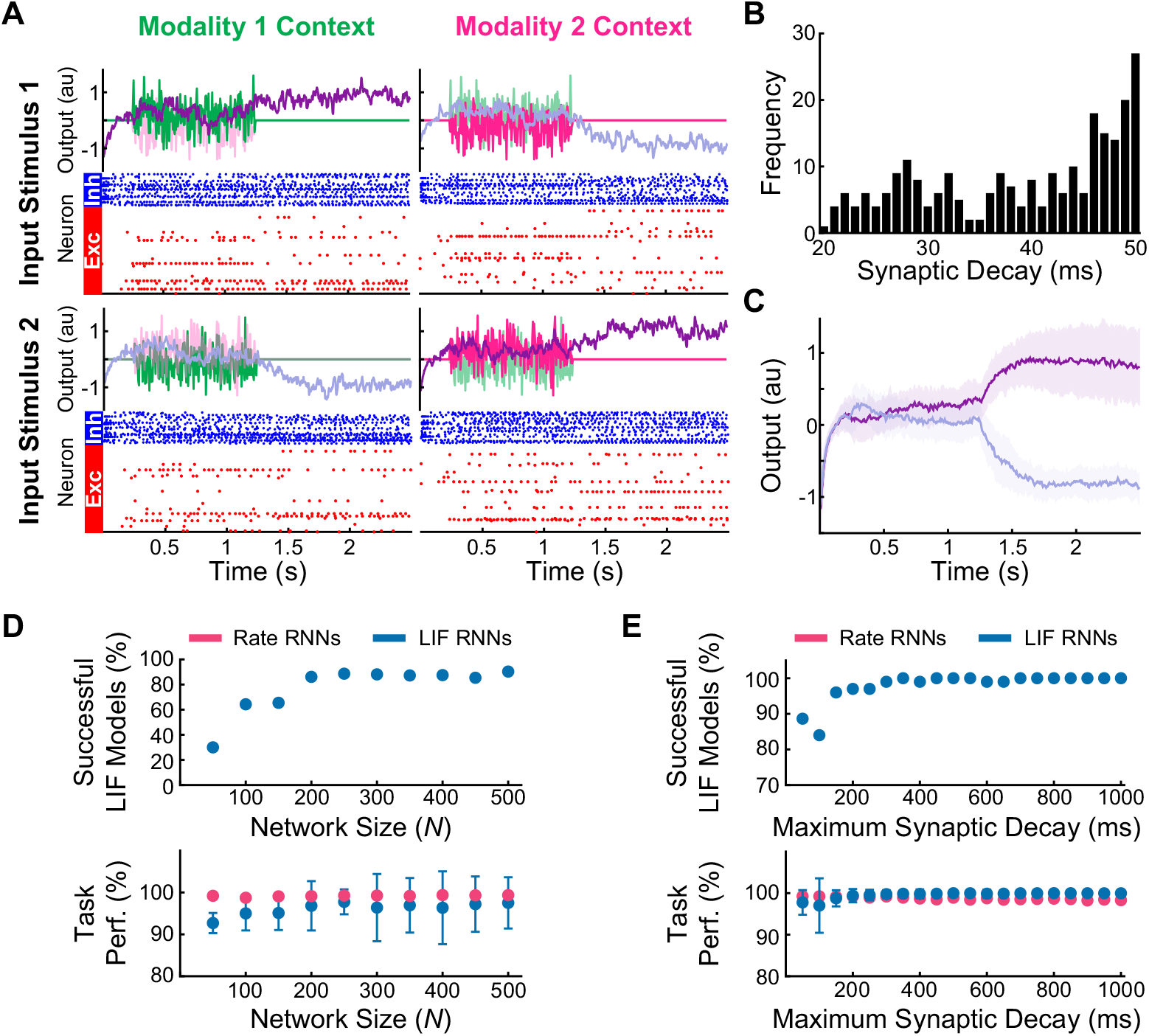
LIF network models constructed to perform the contextual integration task. **A.** Example output responses and spike raster plots from a LIF network model for two different input stimuli (rows) and two contexts (columns). The network contained 250 units (188 excitatory and 62 inhibitory units), and the noisy input signals were scaled by 0.5 vertically for better visualization of the network responses (purple lines). **B.** Distribution of the optimized synaptic decay time constants (***τ***^***d***^) for the example LIF network (Mean ± SD, 38.9 ± 9.3 ms). The time constants were limited to range between 20 ms and 50 ms. **C.** Average output responses of the example LIF network. Mean ± SD network responses across 100 randomly generated trials shown. **D.** Successfully converted LIF networks and their average task performance across different network sizes. The network size was varied from *N* = 10 to 500. The rate RNNs trained in Fig. 3 were used. **E.** Successfully converted LIF networks with *N* = 250 and their average task performance across different maximum synaptic decay constants (varied from 20 ms to 1000 ms).

Similar to the spiking networks constructed for the Go-NoGo task, the LIF RNNs performed the input integration task more accurately as the network size increased (Fig. 4D). Next, the network size was fixed to *N* = 250 and 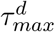 was gradually increased from 20 ms to 1000 ms. For 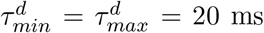, all 100 rate networks failed to learn the task within the first 6000 trials. The conversion from the rate models to the LIF models did not lead to significant loss in task performance for all the other maximum decay constant values considered (Fig. 4E).

### Analysis of the Conversion Method

Previous sections illustrated that our framework for converting rate RNNs to LIF RNNs is robust as long as the network size is not too small (*N* ≥ 200), and the optimal size was *N* = 250 for both tasks. When the network size is too small, it is harder to train rate RNNs and the rate models successfully trained do not reliably translate to spiking networks (Fig. 2D and Fig. 4D). In this section, we further investigate the relationship between rate and LIF RNN models and characterize other parameters crucial for the conversion to be effective.

#### Training synaptic decay time constants

As shown in Fig. 5, training the synaptic decay constants for all the rate units is not required for the conversion to work. Rate RNNs (100 models with different initial conditions) with the synaptic decay time constant fixed to 35 ms (average *τ*^*d*^ value for the networks trained with 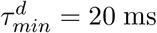 and 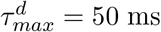) were trained on the Go-NoGo task and converted to LIF RNNs (Fig. 5). The task performance of these LIF networks was not significantly different from the performance of the spiking models with optimized synaptic decay constants bounded between 20 ms and 50 ms. The number of the successful LIF models with the fixed synaptic decay constant was also comparable to the number of the successful LIF models with the tuned decay constants (Fig. 5).

**Fig. 5.**
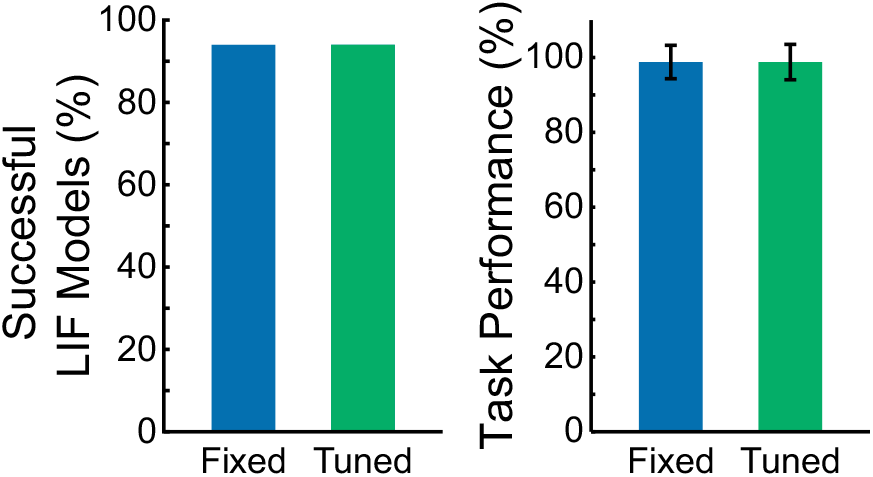
Optimizing synaptic decay constants is not required for conversion of rate RNNs. The Go-NoGo task performance of the LIF RNNs constructed from the rate networks with a fixed synaptic constant (*τ*^*d*^ = 35 ms; blue) was not significantly different from the performance of the LIF RNNs with tuned synaptic decay time constants (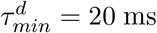, 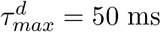; green).

#### Other LIF parameters

We also probed how LIF model parameters affected our framework. More specifically, we focused on the refractory period and synaptic filtering. The LIF models constructed in the previous sections used an absolute refractory period of 2 ms and a double exponential synaptic filter (see Methods). Rate models (*N* = 250 and 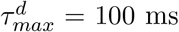) trained on the sensory integration task were converted to LIF networks with different values of the refractory period. As the refractory period became longer, the task performance of the spiking RNNs decreased rapidly (Fig. 6A). When the refractory period was set to 0 ms, the LIF RNNs still performed the integration task with a moderately high average accuracy (92.8 ± 14.3%), but the best task performance was achieved when the refractory period was set to 2 ms (average performance, 97.0 ± 6.6%; Fig. 6A inset).

**Fig. 6.**
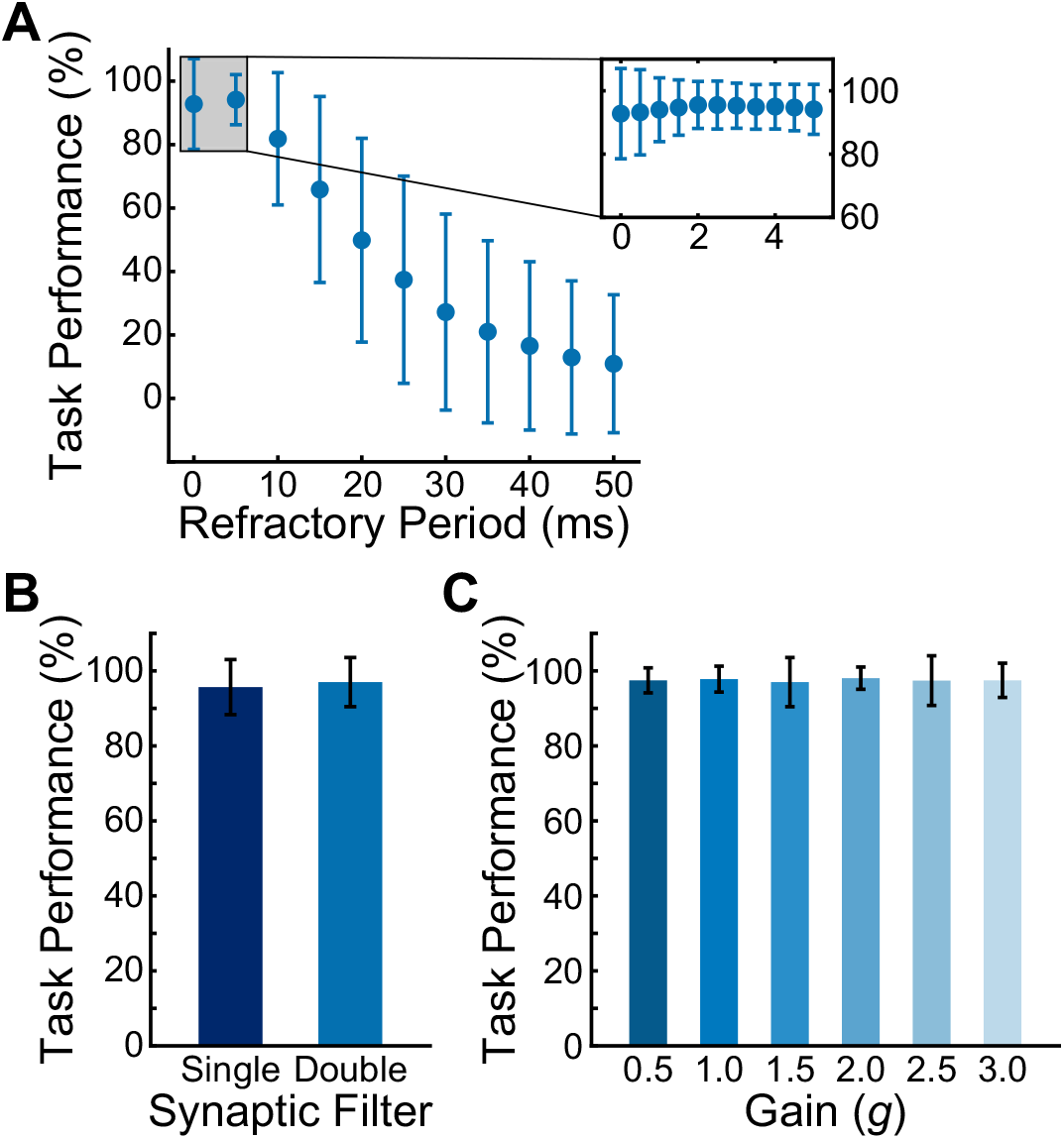
Effects of the refractory period, synaptic filter, and rate RNN connectivity weight initialization. **A.** Average contextual integration task performance of the LIF network models (*N* = 250) with different refractory period values. The refractory period was varied from 0 ms (i.e. no refractory period) to 50 ms. The inset shows the average task performance across finer changes in the refractory period. Mean ± SD shown. **B.** Average contextual integration task performance of the LIF network models (*N* = 250 and refractory period = 2 ms) with the single exponential synaptic filter (dark blue) and the double exponential synaptic filter (light blue). Mean ± SD shown. **C.** Average contextual integration task performance of the LIF network models (*N* = 250, refractory period = 2 ms, and double exponential synaptic filter) with different connectivity gain initializations. Mean ± SD shown.

We also investigated how different synaptic filters influenced the mapping process. We first fixed the refractory period to its optimal value (2 ms) and constructed 100 LIF networks (*N* = 250) for the integration task using a double synaptic filter (see Methods; Fig. 6B light blue). Next, the synaptic filter was changed to the following single exponential filter:

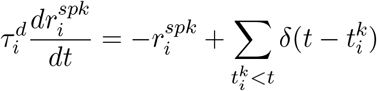

where 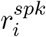 represents the filtered spike train of unit *i* and 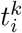 refers to the *k*-th spike emitted by unit *i*. The task performance of the LIF networks with the above single exponential synaptic filter was 95.7 ± 7.3%, and it was not significantly different from the performance of the double exponential synaptic LIF models (97.0 ± 6.6%; Fig. 6B).

#### Initial connectivity weight scaling

We considered the role of the connectivity weight initialization in our framework. In the previous sections, the connectivity weights (*W*^*rate*^) of the rate networks were initialized as random, sparse matrices with zero mean and a standard deviation of 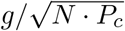, where *g* = 1.5 is the gain term that controls the dynamic regime of the networks and *P*_*c*_ = 0.20 is the initial connectivity probability (see Methods). Previous studies have shown that rate networks operating in a high gain regime (*g >* 1.0) produce chaotic spontaneous trajectories, and this rich dynamics can be harnessed to perform complex computations [6, 11]. By varying the gain term, we determined if highly chaotic initial dynamics were required for successful conversion. We considered six different gain terms ranging from 0.5 to 3.5, and for each gain term, we constructed 100 LIF RNNs (from 100 rate RNNs with random initial conditions; Fig. 6C) to perform the contextual integration task. The LIF models performed the task equally well across all the gain terms considered (no statistical significance detected).

#### Transfer function

One of the most important factors that determines whether rate RNNs can be mapped to LIF RNNs in a one-to-one manner is the nonlinear transfer function used in the rate models. We considered three non-negative transfer functions commonly used in the machine learning field to train rate RNNs on the Go-NoGo task: sigmoid, rectified linear, and softplus functions (Fig. 7A; see SI Appendix). For each transfer function, 100 rate models (*N* = 250 and 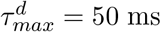) were trained. Although all 300 rate models were trained to perform the task almost perfectly (Fig. 7B), the average task performance and the number of successful LIF RNNs were highest for the rate models trained with the sigmoid transfer function (Fig. 7C). None of the rate models trained with the rectified linear transfer function could be successfully mapped to LIF models, while the spiking networks constructed from the rate models trained with the softplus function were not robust and produced incorrect responses (SI Appendix, Fig. S6).

**Fig. 7.**
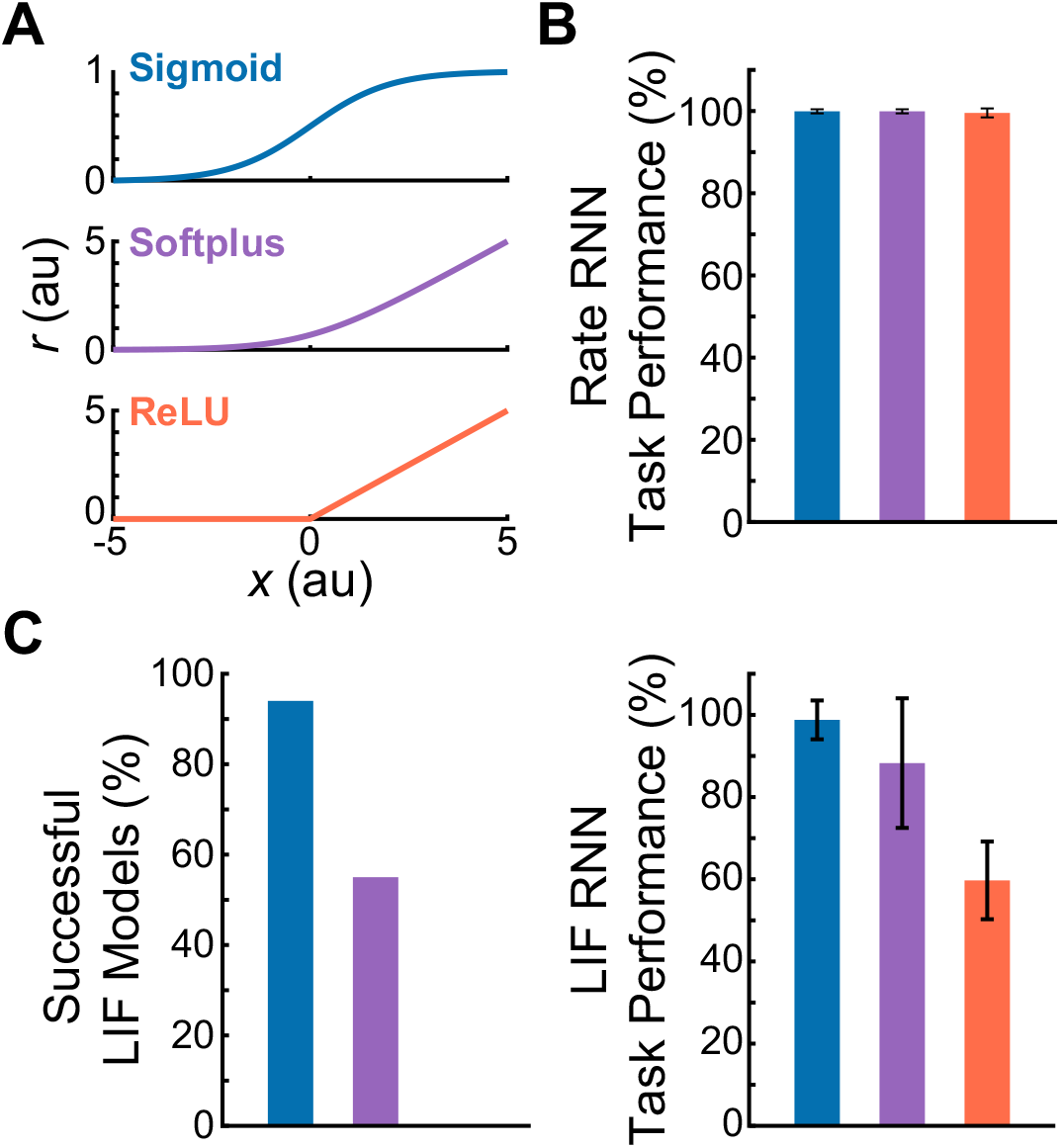
Comparison of the LIF RNNs derived from the rate RNNs trained with three non-negative activation functions. **A.** Three non-negative transfer functions were considered: sigmoid, softplus, and rectified linear (ReLU) functions. **B.** All 300 rate RNNs (100 networks per activation function) were successfully trained to perform the Go-NoGo task. **C.** Of the 100 sigmoid LIF networks constructed, 94 networks successfully performed the task. The conversion rates for the softplus and ReLU LIF models were 55% and 0%, respectively. Mean ± SD task performance: 98.8 ± 4.7% (sigmoid), 88.3 ± 15.8% (softplus), and 59.7 ± 9.5% (ReLU).

## Discussion

In the current study, we presented a simple framework that harnesses the dynamics of trained continuous rate network models to produce functional spiking RNN models. We identified a set of parameters required to directly transform trained rate RNNs to LIF models, thus establishing a one-to-one correspondence between these two model types. Despite of additional spiking-related parameters, surprisingly only a single parameter (i.e. scaling factor) was required for LIF RNN models to closely mimic their counterpart rate models. Furthermore, this framework can flexibly impose functional connectivity constraints and heterogeneous synaptic time constants.

We investigated and characterized the effects of several model parameters on the stability of the transfer learning from rate models to spiking models. The parameters critical for the mapping to be robust included the network size, choice of activation function for training rate RNNs, and a constant factor to scale down the connectivity weights of the trained rate networks. Although the softplus and rectified linear activation functions are popular for training deep neural networks, we demonstrated that the rate networks trained with these functions do not translate robustly to LIF RNNs (Fig. 7). On the other hand, the rate models trained with the sigmoid function were transformed to LIF models with high fidelity.

Another important parameter was the constant scaling factor used to scale *W*^*rate*^ and 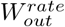 before transferring them to LIF networks. When the scaling factor was set to its optimal value (found via grid search), the LIF units behaved like their counterpart rate units, and the spiking networks performed the tasks the rate RNNs were trained to perform (Fig. 2). Another parameter that affected the reliability of the conversion was the refractory period parameter of the LIF network models. The LIF performance was optimal when the refractory was set to 2 ms (Fig. 6A). Training the synaptic decay time constants, choice of synaptic filter (between single and double exponential filter), and connectivity weight initialization did not affect the mapping procedure (Fig. 5 and Fig. 6B–C).

The type of approach used in this study (i.e. conversion of a rate network to a spiking network) has been previously employed in neuromorphic engineering to construct power-efficient deep spiking networks [31–36]. These studies mainly employed feedforward multi-layer networks or convolutional neural networks aimed to accurately classify input signals or images without placing too much emphasis on biophysical limitations. The overarching goal in these studies was to maximize task performance while minimizing power consumption and computational cost. On the other hand, the main aim of the present study was to construct spiking recurrent network models that abide by important biological constraints in order to relate emerging mechanisms and dynamics to experimentally observed findings. To this end, we have carefully designed our continuous rate RNNs to include several biological features. These include (1) recurrent architectures, (2) sparse connectivity that respects Dale’s principle, and (3) heterogeneous synaptic decay time constants.

For constructing spiking RNNs, recent studies have proposed methods that built on the FORCE method to train spiking RNNs [8, 20–22]. Conceptually, our work is most similar to the work by DePasquale et al. [21]. The method developed by DePasquale et al. [21] also relies on mapping a trained continuous-variable rate RNN to a spiking RNN model. However, the rate RNN model used in their study was designed to provide dynamically rich auxiliary basis functions meant to be distributed to overlapping populations of spiking units. Due to this reason, the relationship between their rate and spiking models is rather complex, and it is not straightforward to impose functional connectivity constraints on their spiking RNN model. An additional procedure was introduced to implement Dale’s principle, but this led to more fragile spiking networks with considerably increased training time [21]. The one-to-one mapping between rate and spiking networks employed in our method solved these problems without sacrificing network stability and computational cost: biophysical constraints that we wanted to incorporate into our spiking model were implemented in our rate network model first and then transferred to the spiking model.

While our framework incorporated the basic yet important biological constraints, there are several features that are also not biologically realistic in our models. The gradient-descent method employed to tune the rate model parameters, including the connectivity weights and the synaptic decay time constants, in a supervised manner is not biologically plausible. Although tuning of the synaptic time constants is not realistic and has not been observed experimentally, previous studies have underscored the importance of the diversity of synaptic time scales both *in silico* and *in vivo* [8, 29, 30]. In addition, other works have validated and uncovered neural mechanisms observed in experimental settings using RNN models trained with backpropagation [7, 13, 37], thus highlighting that a network model can be biologically plausible even if it was constructed using non-biological means. Another limitation of our method is the lack of temporal coding in our LIF models. Since our framework involves rate RNNs that operate in a rate coding scheme, the spiking RNNs that our framework produces also employ rate coding by nature. Previous studies have shown that spike-coding can improve spiking efficiency and enhance network stability [20, 38, 39], and recent studies emphasized the importance of precise spike coordination without modulations in firing rates [40, 41]. Lastly, our framework does not model nonlinear dendritic processes which have been shown to play a significant role in efficient input integration and flexible information processing [22, 42, 43]. Incorporating nonlinear dendritic processes into our platform using the method proposed by Thalmeier et al. [22] will be an interesting next step to further investigate the role of dendritic computation in information processing.

In summary, we provide an easy-to-use platform that converts a continuous recurrent network model with basic biological constraints to a spiking model. The tight relationship between rate and LIF RNN models under certain parameter values suggests that spiking networks could be put together to perform complex tasks traditionally employed to train and study continuous rate networks. Future work needs to focus on why and how such a tight relationship emerges. The framework along with the findings presented in this study lays the groundwork for discovering new principles on how neural circuits solve computational problems with discrete spikes and for constructing more power efficient spiking networks. Extending our platform to incorporate other commonly used neural network architectures could help design biologically plausible deep learning networks that operate at a fraction of the power consumption required for current deep neural networks.

## Acknowledgements

We are grateful to Ben Huh, Gerald Pao, Jason Fleischer, Debha Amatya, Yusi Chen, and Ben Tsuda for helpful discussions and feedback on the manuscript. We also thank Jorge Aldana for assistance with computing resources. This work was funded by the National Institute of Mental Health (F30MH115605-01A1 to R.K.), Harold R. Schwalenberg Medical Scholarship (R.K.), and Burnand-Partridge Foundation Scholarship (R.K.). We also gratefully acknowledge the support of NVIDIA Corporation with the donation of the Quadro P6000 GPU used for this research. The funders had no role in study design, data collection and analysis, decision to publish, or preparation of the manuscript.

## Author contributions

R.K. and T.J.S. designed the study and wrote the manuscript. R.K. and Y.L. performed model analyses and simulations.

## Methods

The implementation of our framework and the codes to generate all the figures in this work are available at https://github.com/rkim35/spikeRNN. The repository also contains implementation of other tasks including autonomous oscillation and exclusive OR (XOR) tasks.

All the trained models used in the present study are available from the corresponding authors upon reasonable request.

### Continuous rate network structure

The continuous rate RNN model contains *N* units recurrently connected to one another. The dynamics of the model is governed by

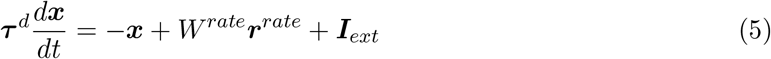

where ***τ***^*d*^ ∈ ℝ^1*×N*^ corresponds to the synaptic decay time constants for the *N* units in the network (see **Training details** on how these are initialized and optimized), ***x***∈ ℝ^1*×N*^ is the synaptic current variable, *W*^*rate*^ ∈ ℝ^*N ×N*^ is the synaptic connectivity matrix, and ***r***^*rate*^ ∈ ℝ^1*×N*^ is the output of the units. The output of each unit, which can be interpreted as the firing rate estimate, is obtained by applying a nonlinear transfer function to the synaptic current variable (***x***) elementwise:

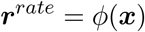

We use a standard logistic sigmoid function for the transfer function to constrain the firing rates to be non-negative:

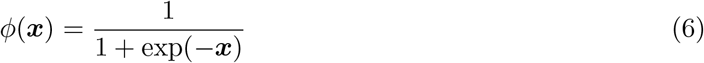

The connectivity weight matrix (*W*^*rate*^) is initialized as a random, sparse matrix drawn from a normal distribution with zero mean and a standard deviation of 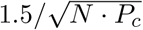 where *P*_*c*_ = 0.20 is the initial connectivity probability.

The external currents (***I***_*ext*_) include task-specific input stimulus signals (see SI Appendix) along with a Gaussian white noise variable:

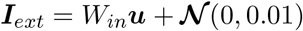

where the time-varying stimulus signals 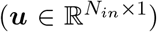 are fed to the network via 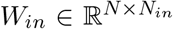, a Gaussian random matrix with zero mean and unit variance. *N*_*in*_ corresponds to the number of input signals associated with a specific task, and 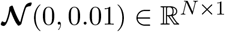 represents a Gaussian random noise with zero mean and variance of 0.01.

The output of the rate RNN at time *t* is computed as a linear readout of the population activity:

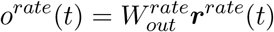

where 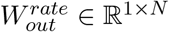 refers to the readout weights.

Eq. (5) is discretized using the first-order Euler approximation method:

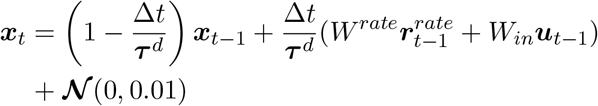

where ∆*t* = 5 ms is the discretization time step size used throughout this study.

### Spiking network structure

For our spiking RNN model, we considered a network of leaky integrate-and-fire (LIF) units governed by

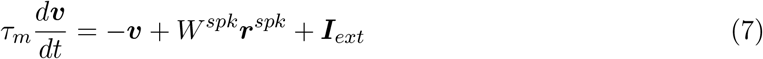

In the above equation, *τ*_*m*_ = 10 ms is the membrane time constant shared by all the LIF units, ***v*** ∈ ℝ^1*×N*^ is the membrane voltage variable, *W*^*spk*^ ∈ ℝ^*N ×N*^ is the recurrent connectivity matrix, and ***r***^*spk*^ ∈ ℝ^1*×N*^ represents the spike trains filtered by a synaptic filter. Throughout the study, the double exponential synaptic filter was used to filter the presynaptic spike trains:

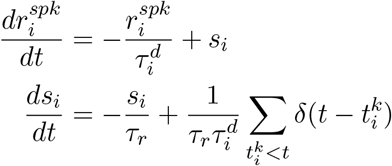

where *τ*_*r*_ = 2 ms and 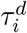 refer to the synaptic rise time and the synaptic decay time for unit *i*, respectively. The synaptic decay time constant values 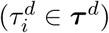 are trained and transferred to our LIF RNN model (see **Training details**). The spike train produced by unit *i* is represented as a sum of Dirac *δ* functions, and 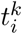 refers to the *k*-th spike emitted by unit *i*.

The external current input (***I***_*ext*_) is similar to the one used in our continuous model (see **Continuous rate network structure**). The only difference is the addition of a constant background current set near the action potential threshold (see below).

The output of our spiking model at time *t* is given by

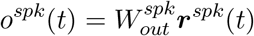

Other LIF model parameters were set to the values used by Nicola et al. [23]. These include the action potential threshold (−40 mV), the reset potential (−65 mV), the absolute refractory period (2 ms), and the constant bias current (−40 pA). The parameter values for the LIF and the quadratic integrate-and-fire (QIF) models are listed in SI Appendix, Table S1.

### Training details

In this study, we only considered supervised learning tasks. A task-specific target signal (***z***) is used along with the rate RNN output (***o***^*rate*^) to define the loss function 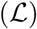, which our rate RNN model is trained to minimize. Throughout the study, we used the root mean squared error (RMSE) defined as

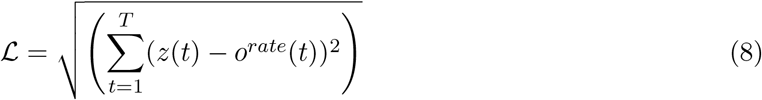

where *T* is the total number of time points in a single trial.

In order to train the rate model to minimize the above loss function (Eq. 8), we employed Adaptive Moment Estimation (ADAM) stochastic gradient descent algorithm. The learning rate was set to 0.01, and the TensorFlow default values were used for the first and second moment decay rates. The gradient descent method was used to optimize the following parameters in the rate model: synaptic decay time constants (***τ***^*d*^), recurrent connectivity matrix (*W*^*rate*^), and readout weights 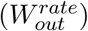.

Here we describe the method to train synaptic decay time constants (***τ***^*d*^) using backpropagation. First, the time constants are initialized with random values within the specified range:

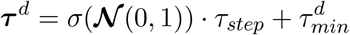

where *σ*(⋅) is the sigmoid function (identical to Eq. 6) used to constrain the time constants to be non-negative. The time constant values are also bounded by the minimum 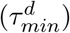 and themaximum 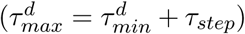 values. The error computed from the loss function (Eq. 8) is then backpropagated to update the time constants at each iteration:

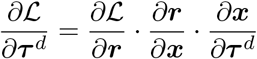

The method proposed by Song et al. [13] was used to impose Dale’s principle and create separate excitatory and inhibitory populations. Briefly, the recurrent connectivity matrix (*W*^*rate*^) in the rate model is parametrized by

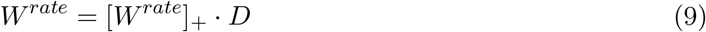

where the rectified linear operation ([⋅]_+_) is applied to the connectivity matrix at each update step. The diagonal matrix (*D* ∈ −^*N* ×*N*^) contains +1’s for excitatory units and −1’s for inhibitory units in the network. Each unit in the network is randomly assigned to one group (excitatory or inhibitory) before training, and the assignment does not change during training (i.e. *D* stays fixed).

To impose specific connectivity patterns, we apply a binary mask (*M* ∈ ℝ^*N ×N*^) to Eq. 9:

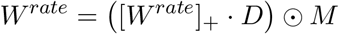

where ⊙ refers to the Hadamard operation (elementwise multiplication). Similar to the diagonal matrix (*D*), the mask matrix stays fixed throughout training. For example, the following mask matrix can be used to create a subgroup of inhibitory units (Group A) that do not receive synaptic inputs from the rest of the inhibitory units (Group B) in the network (Fig. S3):

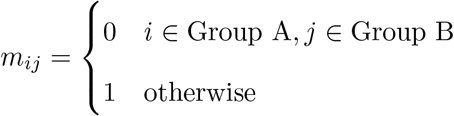

where *m*_*ij*_ ∈ *M* establishes (if *m*_*ij*_ = 1) or removes (if *m*_*ij*_ = 0) the connection from unit *j* to unit *i*.

### Transfer learning from a rate model to a spiking model

In this section, we describe the method that we developed to perform transfer learning from a trained rate model to a LIF model. Once the rate RNN model is trained using the gradient descent method, the rate model parameters are transferred to a LIF network in a one-to-one manner. First, the LIF network is initialized to have the same topology as the trained rate RNN. Next, the input weight matrix (*W*_*in*_) and the synaptic decay time constants (***τ***^*d*^) are transferred to the spiking RNN without any modification. Lastly, the recurrent connectivity matrix (*W*^*rate*^) and the readout weights 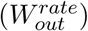 are scaled by a constant number, *λ*, and transferred to the spiking network.

If the recurrent connectivity weights from the trained rate model are transferred to a spiking network without any changes, the spiking model produces largely fluctuating signals (as illustrated in Fig. 2B), because the LIF firing rates are significantly larger than 1 (whereas the firing rates of the rate model are constrained to range between zero and one by the sigmoid transfer function).

To place the spiking RNN in the similar dynamic regime as the rate network, we first assume a linear relationship between the rate model connectivity weights and the spike model weights:

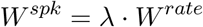

Using the above assumption, the synaptic drive (*d*) that unit *i* in the LIF RNN receives can be expressed as

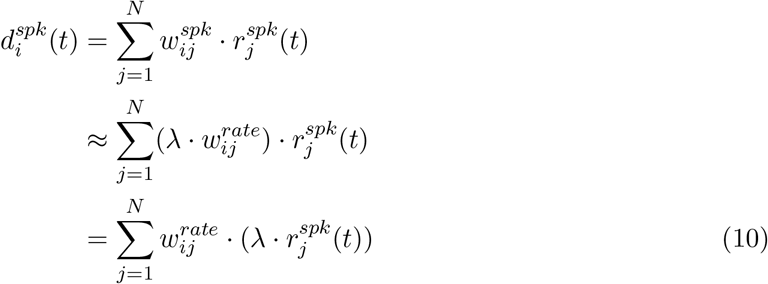

where 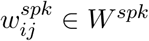 is the synaptic weight from unit *j* to unit *i*.

Similarly, unit *i* in the rate RNN model receives the following synaptic drive at time *t*:

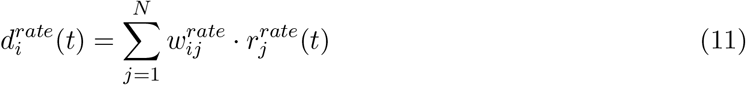

If we set the above two synaptic drives (Eq. 10 and Eq. 11) equal to each other, we have:

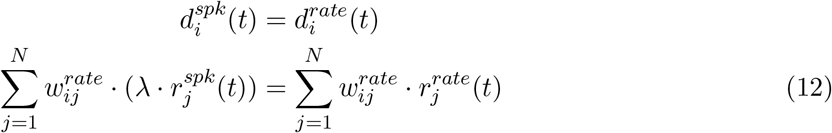

Generalizing Eq. 12 to all the units in the network, we have

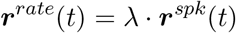

Therefore, if there exists a constant factor (*λ*) that can account for the firing rate scale difference between the rate and the spiking models, the connectivity weights from the rate model (*W*^*rate*^) can be scaled by the factor and transferred to the spiking model.

The readout weights from the rate model 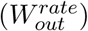 are also scaled by the same constant factor (*λ*) to have the spiking network produce output signals similar to the ones from the trained rate model:

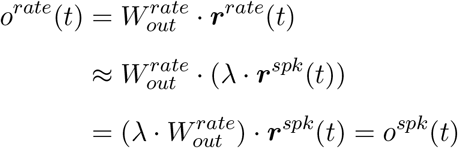

In order to find the optimal scaling factor, we developed a simple grid search algorithm. For a given range of values for 1*/λ* (ranged from 20 to 75 with a step size of 5), the algorithm finds the optimal value that maximizes the task performance.

### Implementation of computational tasks and figure details

In this section, we describe the details of the parameters and methods used to generate all the main figures in the present study.

*Fig. 1.* A rate RNN of *N* = 200 units (169 excitatory and 31 inhibitory units) was trained to perform the Go-NoGo task for Fig. 1B. Each trial lasted for 1000 ms (200 time steps with 5 ms step size). The minimum and the maximum synaptic decay time constants were set to 20 ms and 50 ms, respectively. An input stimulus with a pulse 125 ms in duration was given for a Go trial, while no input stimulus was given for a NoGo trial. The network was trained to produce an output signal approaching +1 after the stimulus offset for a Go trial. For a NoGo trial, the network was trained to maintain its output at zero. A trial was considered correct if the maximum output signal during the response window was above 0.7 for the Go trial type. For a NoGo trial, if the maximum response value was less than 0.3, the trial was considered correct. For training, 6000 trials were randomly generated, and the model performance was evaluated after every 100 trials. Training was terminated when the loss function value fell below 7 and the task performance reached at least 95%. The termination criteria were usually met at or before 2000 trials for this task.

For Fig. 1C, rate RNNs with 9 different sizes (*N* = 10, 50, 100, 150, 200, 250, 300, 350, 400) were trained. For each network size, 100 rate RNNs with random initial conditions were trained on the Go-NoGo task.

*Fig. 2.* The rate RNN trained in Fig. 1B was converted to a LIF RNN using different scaling factor (*λ*) values for Fig. 2B. The double exponential synaptic filter was used, and the gain term (*g*) for the rate RNN initialization was set to 1.5. The LIF parameters listed in Table S1 were used for all the LIF network models constructed in Fig. 2.

*Fig. 3.* Rate RNNs with 11 different network sizes (*N* = 10, 50, 100, 150, 200, 250, 300, 350, 400, 450, 500) were trained on the contextual integration task. For each network size, 100 rate RNNs with random initial conditions were trained.

For the task design, the input matrix (***u*** ∈ ℝ^4*×*500^) contained four stimuli channels across time (500 time steps with 5 ms step size). The first two channels corresponded to the modality 1 and modality 2 noisy input signals. These signals were modeled as white-noise signals (sampled from the standard normal distribution) with constant offset terms. The sign of the offset term modeled the evidence toward (+) or (−) choices, while the magnitude of the offset determined the strength of the evidence. The noisy signals were only present during the stimulus window (250 ms – 1250 ms). The last two channels of ***u*** represented the modality 1 and the modality 2 context signals. For instance, the third channel of ***u*** is set to one and the fourth channel is set to zero throughout the trial duration to model Modality 1 context.

For each trial used to train the rate model, the offset values for the two modality input signals were randomly set to −0.5 or +0.5. The context signals were randomly set such that either modality 1 (third input channel is set to 1) or modality 2 (fourth input channel is set to 1) was cued for each trial. If the offset term of the cued modality was +0.5 (or −0.5) for a given trial, the network was instructed to produce an output signal approaching +1 (or −1) after the stimulus window. The model performance was assessed after every 100 training trials, and the training termination conditions were same as the ones used for Fig. 1.

*Fig. 4.* A network of *N* = 250 LIF units (188 excitatory and 62 inhibitory units) were constructed from a rate RNN model trained to perform the context-dependent input integration task for Fig. 4A. The scaling factor (*λ*) was set to 1/60. The double exponential synaptic filter was used, and the gain term (*g*) for the rate RNN initialization was set to 1.5. The LIF parameters listed in Table S1 were used for all the LIF network models constructed in Fig. 4.

*Fig. 5.* Rate RNNs (*N* = 250) were trained on the Go-NoGo task with and without optimizing the synaptic decay time constants (***τ***^***d***^). For each condition, 100 rate RNNs were trained. For the fixed synaptic decay constant condition, *τ*^*d*^ was fixed to 35 ms. For the tuned synaptic decay condition, 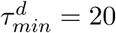 and 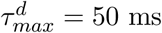.

*Fig. 6.* For Fig. 6A, all 100 rate RNNs (*N* = 250, 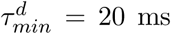, 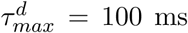) trained in Fig. 4E were converted to LIF RNNs with different values of the refractory period. The following 20 refractory period values were considered: 0, 0.5, 1.0, 1.5, 2.0, 2.5, 3.0, 3.5, 4.0, 4.5, 5.0, 10, 15, 20, 25, 30, 35, 40, 45, 50 ms.

*Fig. 7.* The following softplus function was used:

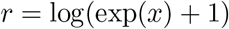

For the networks trained with the softplus and ReLU activation functions, the following range of values for 1*/λ* was used for the grid search: 4 to 26 with a step size of 2.

**Quadratic integrate-and-fire model.** For the quadratic integrate-and-fire (QIF) model (Fig. S7), we considered a network of units governed by

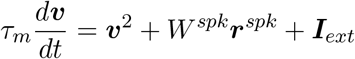

The definitions of the variables are identical to the ones used for the LIF network model.

---

**Supplementary Fig. 1.**
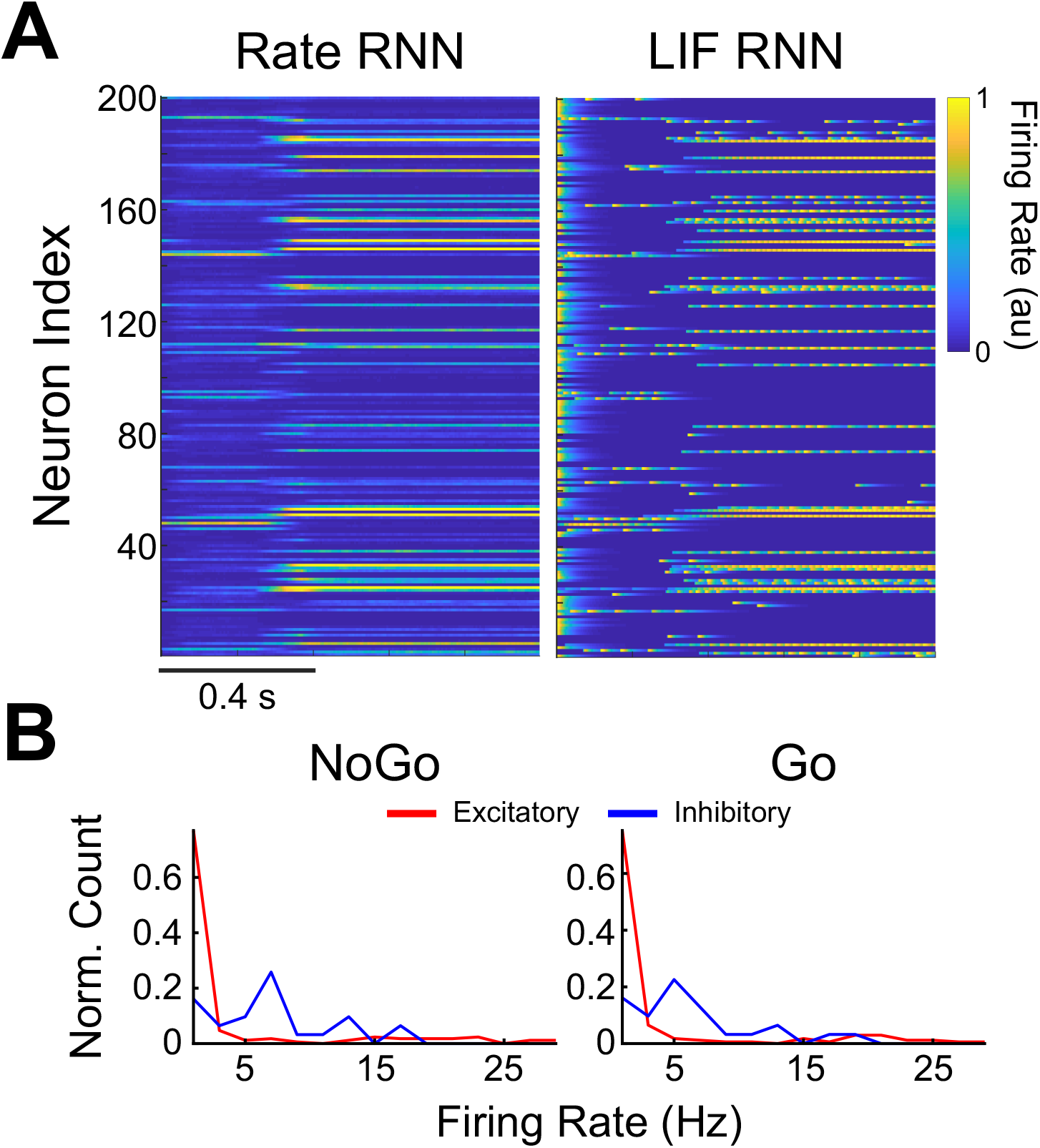
Comparison of the time-varying rates of the continuous-variable rate units and the LIF units. **A.** A single Go trial was used to extract the rates from the rate RNN trained in Fig. 1B. The firing rates of the LIF RNN constructed using the optimal scaling factor (*λ* = 1/25) are shown on the right. The firing rates of the LIF units were normalized to range from 0 to 1 for comparison. **B.** Distribution of the firing rates for a NoGo trial (left) and a Go trial (right).

**Supplementary Fig. 2.**
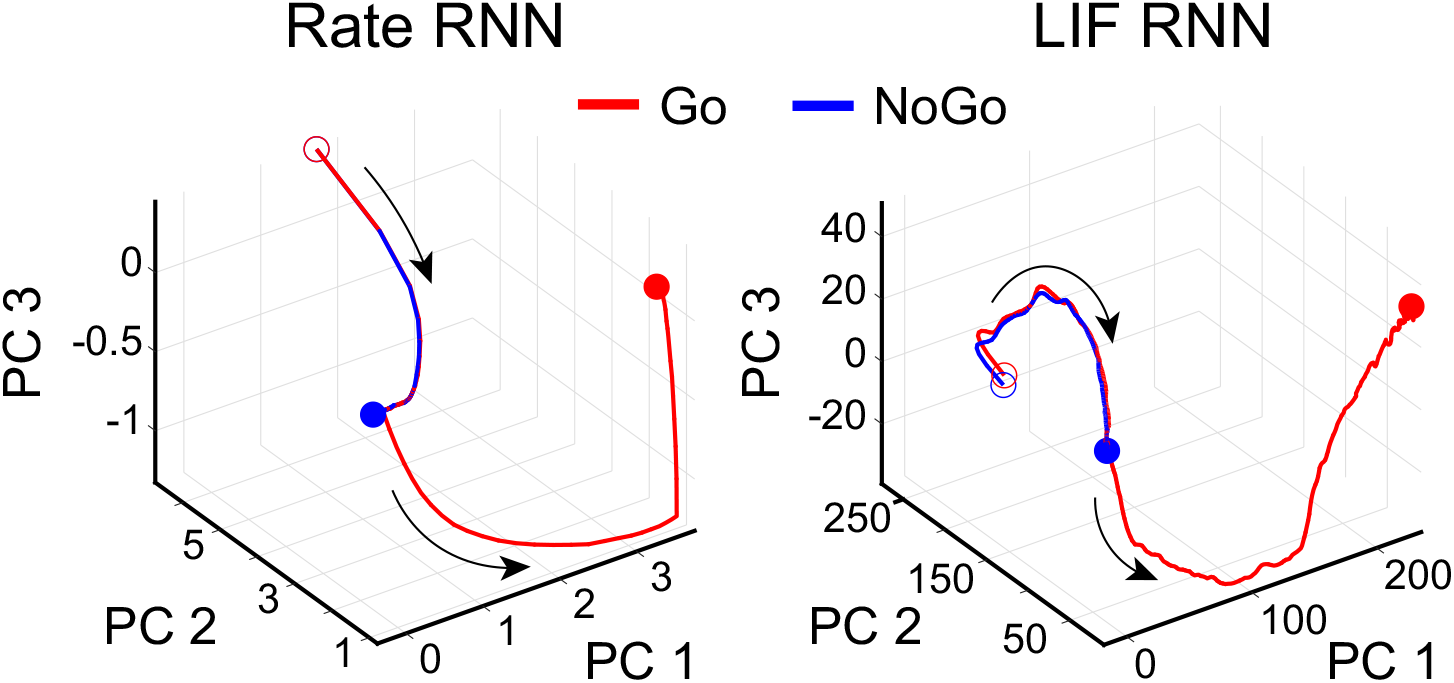
Comparison of the top three PCs extracted from the network activities of the rate and LIF RNNs trained to perform the Go-NoGo task. Principal component analysis (PCA) was performed on the firing rates derived from a rate RNN and a LIF RNN trained to perform the Go-NoGo task. The rate RNN contained 200 units (169 excitatory and 31 inhibitory units), and the LIF model was constructed from the rate model. The firing rates from 50 Go trials and 50 NoGo trials were obtained from the two RNN models. For both models, the top three principal components (PCs) captured 99% of the variance. Red and blue empty circles indicate the trial onset for the Go and the NoGo trials, respectively. Red and blue filled circles represent the end of the trial for the Go and the NoGo trials, respectively.

**Supplementary Fig. 3.**
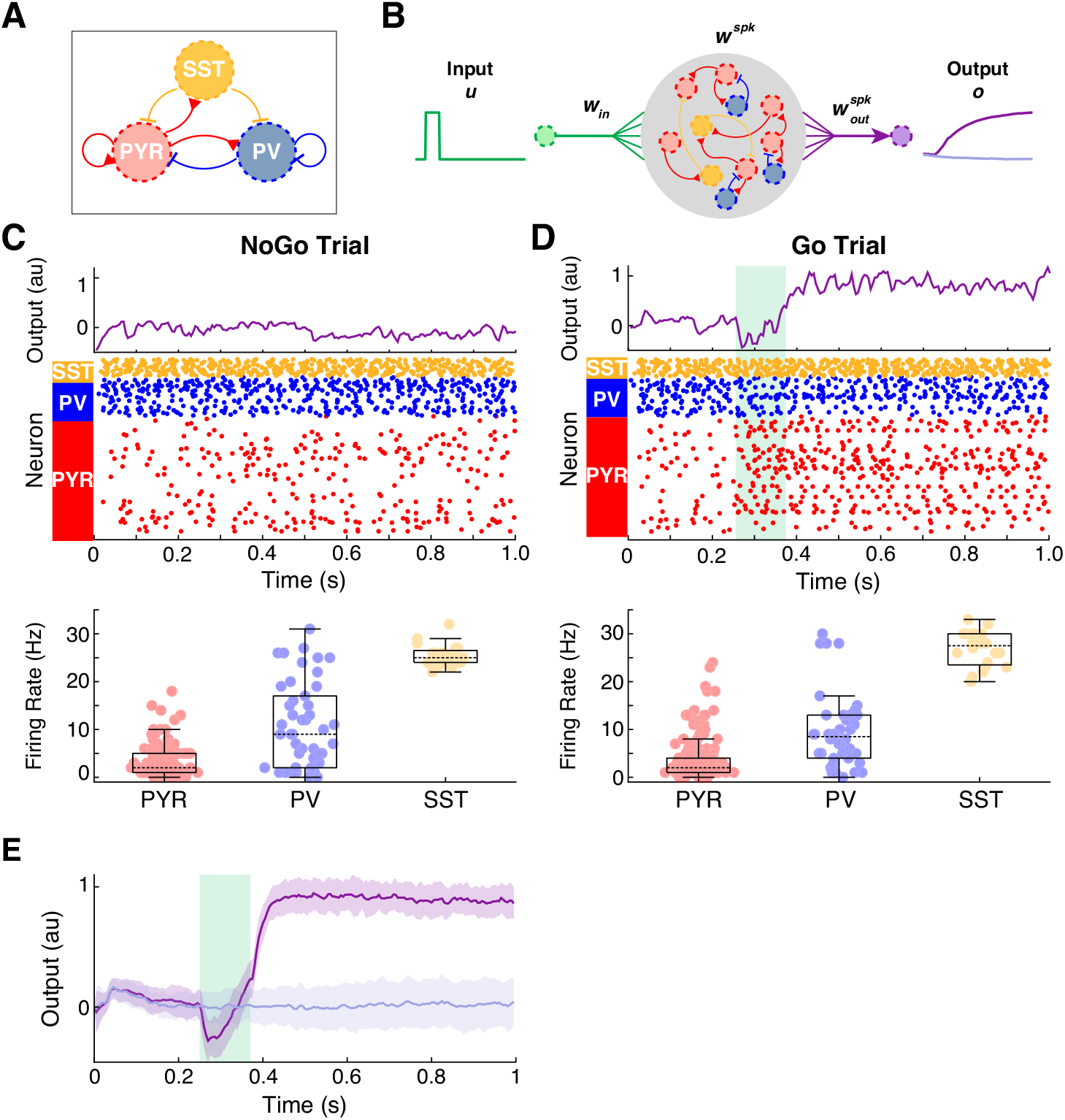
Incorporation of additional functional connectivity constraints. **A.** Common cortical microcircuit motif where somatostatin-expressing interneurons (SST; yellow circle) inhibit both pyramidal (PYR; red circle) and parvalbumin-expressing (PV; blue circle) neurons. **B.** Schematic illustrating the incorporation of the connectivity motif shown in **A** into a LIF network model. The connectivity pattern was imposed during training of a rate network model (*N* = 200) to perform the Go-NoGo task. There were 134 PYR, 46 PV, and 20 SST units. A spiking model was constructed using the trained rate model with *λ* = 1/50. **C.** Example output response and spikes from the LIF network model for a single NoGo trial. Mean ± SD firing rate for each population is also shown (PYR, 3.08 ± 3.29 Hz; PV, 10.80 ± 8.94 Hz; SST, 25.50 ± 2.33 Hz). **D.** Example output response and spikes from the LIF network model for a single Go trial. Mean ± SD firing rate for each population is also shown (PYR, 4.72 ± 5.89 Hz; PV, 9.30 ± 8.16 Hz; SST, 27.05 ± 3.98 Hz). Box plot central lines, median; bottom and top edges, lower and upper quartiles. **E.** LIF network model performance on 50 NoGo trials (light purple) and 50 Go trials (dark purple). Mean ± SD shown.

**Supplementary Fig. 4.**
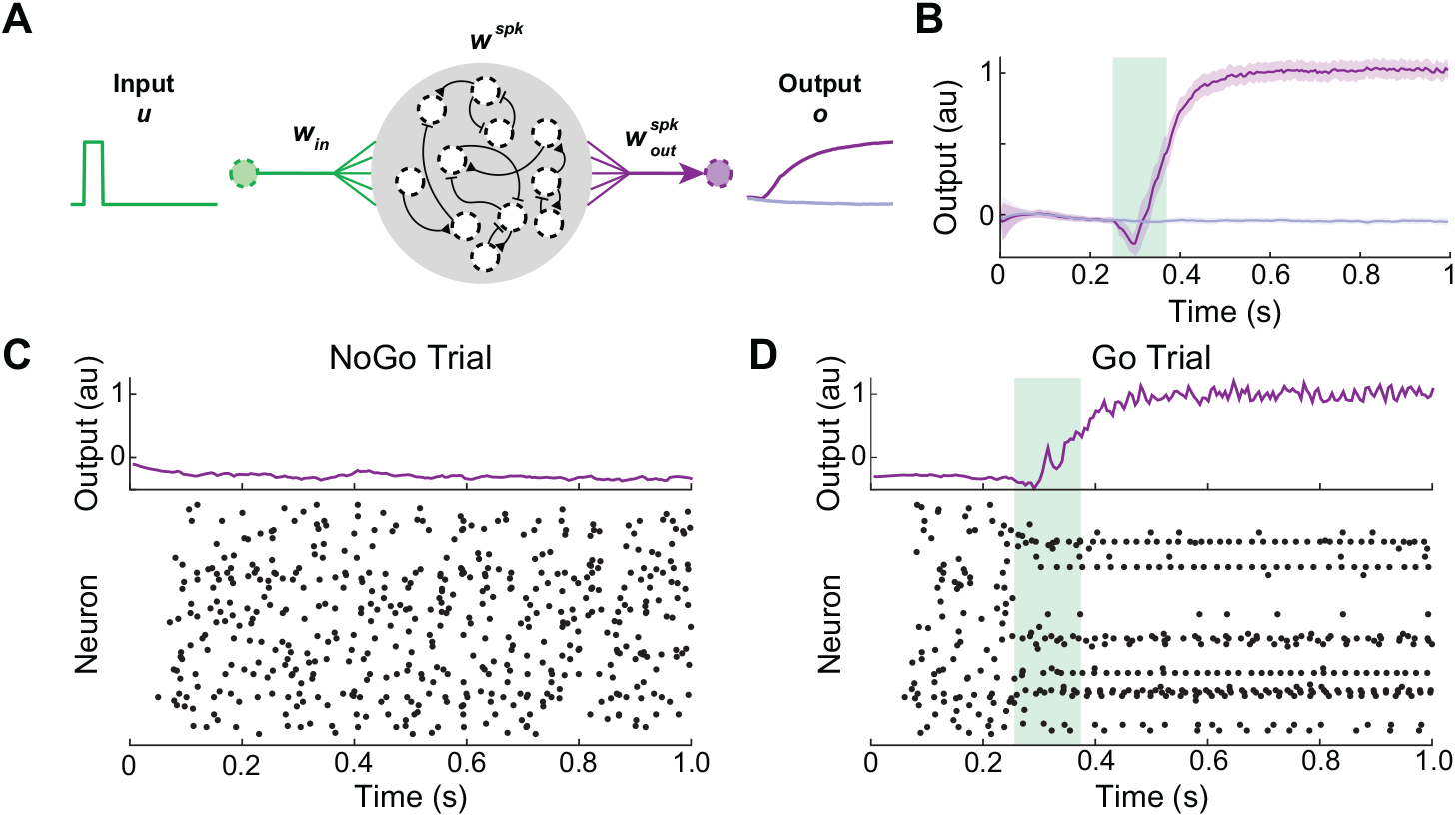
Dale’s principle constraint can be relaxed. **A.** Schematic diagram showing a LIF network model without Dale’s principle. A rate RNN model (*N* = 200) without Dale’s principle was first trained to perform the Go-NoGo task. The scaling factor (*λ*) was set to 1/50. Note that each unit (black dotted circles) can exert both excitatory and inhibitory effects. **B.** LIF network model performance on 50 NoGo trials (light purple) and 50 Go trials (dark purple). Mean ± SD shown. **C.** Example output response (top) and spikes (bottom) from the LIF network model for a single NoGo trial. **D.** Example output response (top) and spikes (bottom) from the LIF network model for a single Go trial.

**Supplementary Fig. 5.**
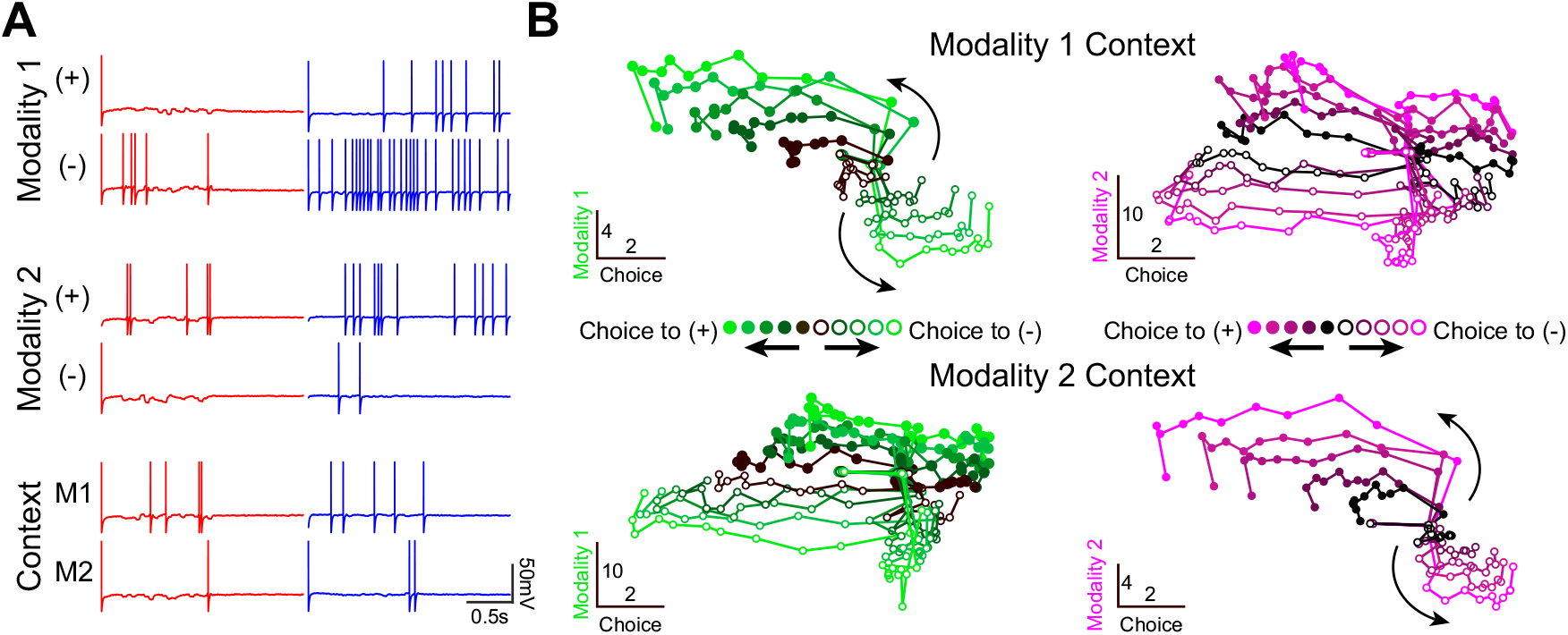
The LIF network model employs mixed representations of the task variables. **A.** Mixed representation of the task variables at the level of single units from a LIF network (*N* = 400; 299 excitatory and 101 inhibitory units; 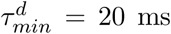 and 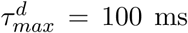). An excitatory unit (red) and an inhibitory unit (blue) with mixed representation of three task variables (modality 1, modality 2, and context) are shown as examples. The excitatory neuron preferred modality 1 input signals with negative offset values, modality 2 signals with positive offset values, and modality 1 context (left column). The inhibitory neuron also exhibited similar biases (right column). **B.** Average population responses projected to a low dimensional state space. The targeted dimensionality reduction technique (developed in [7]) was used to project the population activities to the state space spanned by the task-related axes. For the modality 1 context (top row), the population responses from the trials with various modality 1 offset values were projected to the choice and modality 1 axes (left). The same trials were sorted by the irrelevant modality (modality 2) and shown on the right. Similar conventions used for the modality 2 context (bottom row). The offset magnitude (i.e. amount of evidence toward “+” or “-” choice) increases from dark to light. Filled and empty circles correspond to “+” choice and “-” choice trials, respectively.

**Supplementary Fig. 6.**
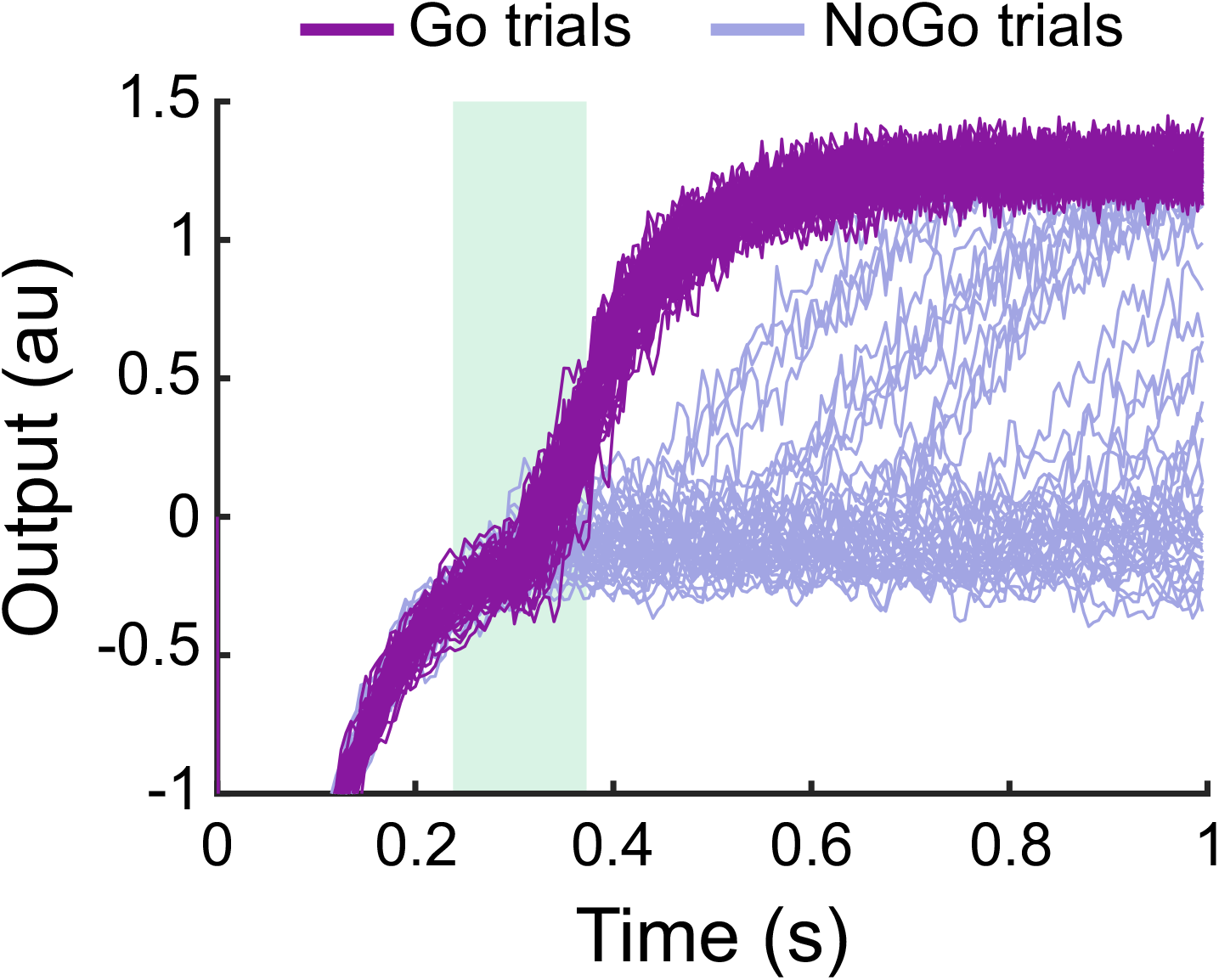
Example output responses from a softplus LIF RNN constructed to perform the Go-NoGo task. Individual output responses from 50 Go trials (dark purple) and 50 NoGo trials (light purple) are shown. The optimal scaling factor was 1/10, and the performance of the model was 78%.

**Supplementary Fig. 7.**
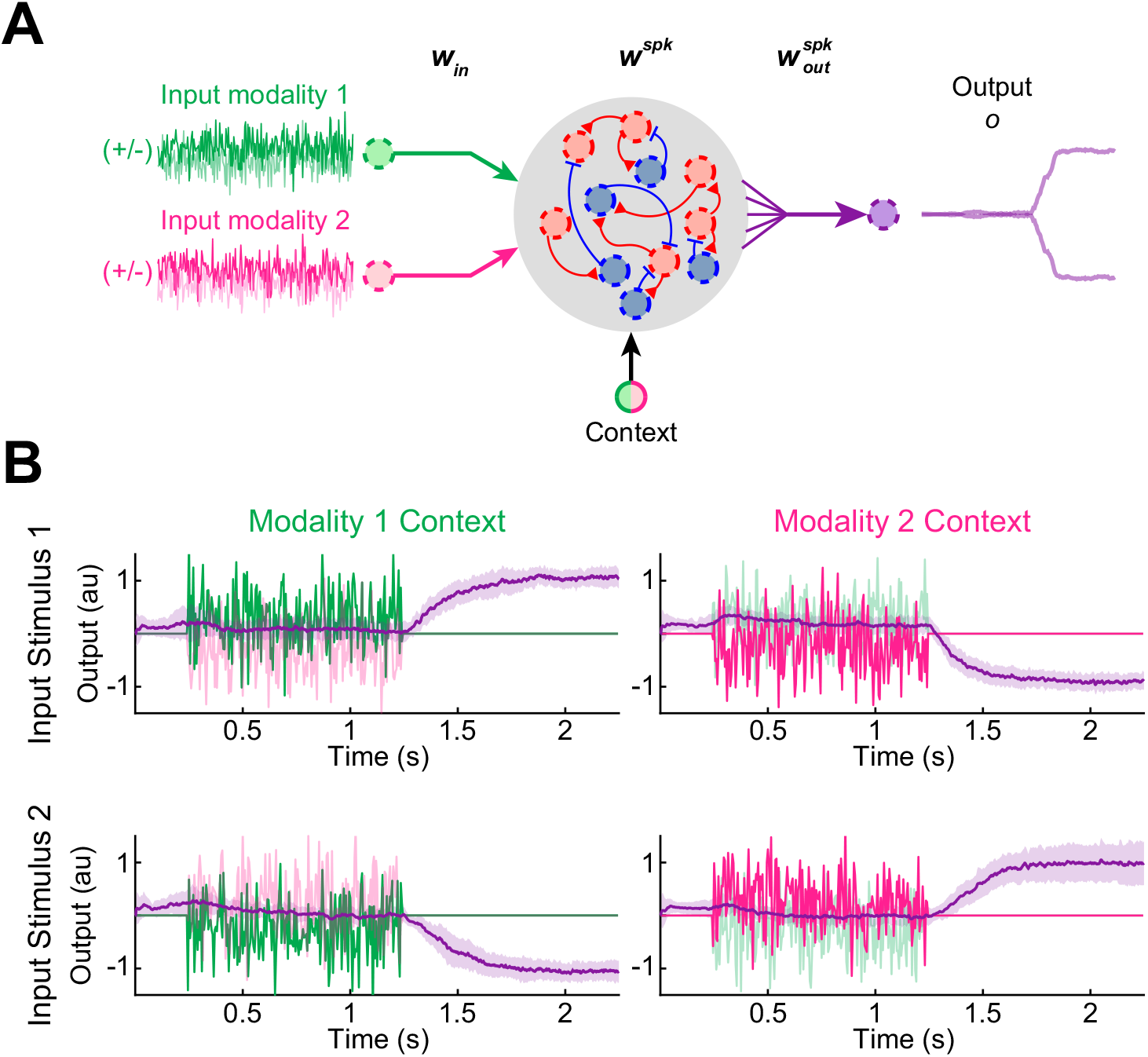
Quadratic integrate-and-fire (QIF) model constructed to perform the context-dependent input integration task. **A.** The task paradigm and the trained rate network model used for Fig. S5 were employed to build a QIF model. The QIF model parameter values are listed in Table S1. **B.** The QIF model successfully performed the task by integrating cued modality input signals. Example noisy input signals (scaled by 0.5 vertically for visualization; green and magenta lines) from a single trial are shown. Mean ± SD response signals (purple lines) across 50 trials for each trial type.

## Supplementary Notes

For the quadratic integrate-and-fire (QIF) model (Supplementary Fig. 7), we considered a network of units governed by

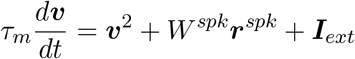

The definitions of the variables are identical to the ones used for the LIF network model.

---

**Supplementary Table 1.** Parameter values used to construct LIF and QIF networks.

|  | LIF | QIF |
| --- | --- | --- |
| Membrane time constant ($\tau_m$) | 10 ms | 10 ms |
| Absolute refractory period | 2 ms | 2 ms |
| Synaptic rise time ($\tau_r$) | 2 ms | 2 ms |
| Constant bias current | -40 pA | 0 pA |
| Spike threshold | -40 mV | 30 mV |
| Spike reset voltage | -65 mV | -65 mV |

## References

[1] Goldman-Rakic, P. Cellular basis of working memory. Neuron. 14, 477–485. (1995).

[2] Felsen, G., song Shen, Y., Yao, H., Spor, G., Li, C. & Dan, Y. Dynamic modification of cortical orientation tuning mediated by recurrent connections. Neuron. 36, 945–954. (2002).

[3] Wang, X.-J. Decision making in recurrent neuronal circuits. Neuron. 60, 215–234. (2008).

[4] Sompolinsky, H., Crisanti, A. & Sommers, H. J. Chaos in random neural networks. Phys. Rev. Lett.. 61, 259–262 (1988).

[5] Sussillo, D. & Abbott, L. Generating coherent patterns of activity from chaotic neural networks. Neuron. 63, 544–557 (2009).

[6] Laje, R. & Buonomano, D. V. Robust timing and motor patterns by taming chaos in recurrent neural networks. Nature Neuroscience. 16, 925–933 (2013).

[7] Mante, V., Sussillo, D., Shenoy, K. V. & Newsome, W. T. Context-dependent computation by recurrent dynamics in prefrontal cortex. Nature. 503, 78–84 (2013).

[8] Kim, C. M. & Chow, C. C. Learning recurrent dynamics in spiking networks. eLife. 7, e37124 (2018).

[9] Mastrogiuseppe, F. & Ostojic, S. Linking connectivity, dynamics, and computations in low-rank recurrent neural networks. Neuron. 99, 609–623 (2018).

[10] Enel, P., Procyk, E., Quilodran, R. & Dominey, P. F. Reservoir computing properties of neural dynamics in prefrontal cortex. PLOS Computational Biology. 12, e1004967 (2016).

[11] Rajan, K., Harvey, C. D. & Tank, D. W. Recurrent network models of sequence generation and memory. Neuron. 90, 128–142 (2016).

[12] Barak, O., Sussillo, D., Romo, R., Tsodyks, M. & Abbott, L. From fixed points to chaos: Three models of delayed discrimination. Progress in Neurobiology. 103, 214–222 (2013).

[13] Song, H. F., Yang, G. R. & Wang, X.-J. Training excitatory-inhibitory recurrent neural networks for cognitive tasks: A simple and flexible framework.. PLOS Computational Biology. 12, e1004792 (2016).

[14] Song, H. F., Yang, G. R. & Wang, X.-J. Reward-based training of recurrent neural networks for cognitive and value-based tasks. eLife. 6, e21492 (2017).

[15] Miconi, T. Biologically plausible learning in recurrent neural networks reproduces neural dynamics observed during cognitive tasks. eLife. 6, e20899 (2017).

[16] Wang, J. X., Kurth-Nelson, Z., Kumaran, D., Tirumala, D., Soyer, H., Leibo, J. Z., Hassabis, D. & Botvinick, M. Prefrontal cortex as a meta-reinforcement learning system. Nature Neuroscience. 21, 860–868. (2018).

[17] Zhang, Z., Cheng, Z., Lin, Z., Nie, C. & Yang, T. A neural network model for the orbitofrontal cortex and task space acquisition during reinforcement learning. PLOS Computational Biology. 14, 1–24. (2018).

[18] Huh, D. & Sejnowski, T. J. Gradient descent for spiking neural networks. In Bengio, S., Wallach, H., Larochelle, H., Grauman, K., Cesa-Bianchi, N. & Garnett, R., editors, Advances in Neural Information Processing Systems 31. pages 1433–1443 (2018).

[19] Lee, J. H., Delbruck, T. & Pfeiffer, M. Training deep spiking neural networks using backpropagation. Frontiers in Neuroscience. 10, 508 (2016).

[20] Abbott, L. F., DePasquale, B. & Memmesheimer, R.-M. Building functional networks of spiking model neurons. Nature Neuroscience. 19, 350–355 (2016).

[21] DePasquale, B., Churchland, M. M. & Abbott, L. F. Using firing-rate dynamics to train recurrent networks of spiking model neurons. Preprint at arXiv https://arxiv.org/abs/1601.07620 (2016).

[22] Thalmeier, D., Uhlmann, M., Kappen, H. J. & Memmesheimer, R.-M. Learning universal computations with spikes. PLOS Computational Biology. 12, e1004895 (2016).

[23] Nicola, W. & Clopath, C. Supervised learning in spiking neural networks with force training. Nature Communications. 8, 2208 (2017).

[24] Werbos, P. J. Backpropagation through time: what it does and how to do it. Proceedings of the IEEE. 78, 1550–1560 (1990).

[25] Martens, J. & Sutskever, I. Learning recurrent neural networks with hessian-free optimization. In Proceedings of the 28th International Conference on International Conference on Machine Learning. ICML’11. pages 1033–1040. USA. (2011). Omnipress.

[26] Pascanu, R., Mikolov, T. & Bengio, Y. On the difficulty of training recurrent neural networks. In Proceedings of the 30th International Conference on International Conference on Machine Learning - Volume 28. ICML’13. pages III–1310–III–1318. (2013).

[27] Bengio, Y., Boulanger-Lewandowski, N. & Pascanu, R. Advances in optimizing recurrent networks. In 2013 IEEE International Conference on Acoustics, Speech and Signal Processing. pages 8624–8628. (2013).

[28] Stokes, M. G., Kusunoki, M., Sigala, N., Nili, H., Gaffan, D. & Duncan, J. Dynamic coding for cognitive control in prefrontal cortex. Neuron. 78, 364–375. (2013).

[29] Wasmuht, D. F., Spaak, E., Buschman, T. J., Miller, E. K. & Stokes, M. G. Intrinsic neuronal dynamics predict distinct functional roles during working memory. Nature Communications. 9, 3499 (2018).

[30] Cavanagh, S. E., Towers, J. P., Wallis, J. D., Hunt, L. T. & Kennerley, S. W. Reconciling persistent and dynamic hypotheses of working memory coding in prefrontal cortex. Nature Communications. 9. (2018).

[31] Cao, Y., Chen, Y. & Khosla, D. Spiking deep convolutional neural networks for energy-efficient object recognition. Int. J. Comput. Vision. 113, 54–66. (2015).

[32] Diehl, P. U., Neil, D., Binas, J., Cook, M., Liu, S. & Pfeiffer, M. Fast-classifying, high-accuracy spiking deep networks through weight and threshold balancing. In 2015 International Joint Conference on Neural Networks (IJCNN). pages 1–8. (2015).

[33] Diehl, P. U., Zarrella, G., Cassidy, A., Pedroni, B. U. & Neftci, E. Conversion of artificial recurrent neural networks to spiking neural networks for low-power neuromorphic hardware. In 2016 IEEE International Conference on Rebooting Computing (ICRC). pages 1–8. (2016).

[34] Hunsberger, E. & Eliasmith, C. Training spiking deep networks for neuromorphic hardware. CoRR. abs/1611.05141. (2016).

[35] Rueckauer, B., Lungu, I.-A., Hu, Y. & Pfeiffer, M. Theory and tools for the conversion of analog to spiking convolutional neural networks. (2016).

[36] Sengupta, A., Ye, Y., Wang, R., Liu, C. & Roy, K. Going deeper in spiking neural networks: Vgg and residual architectures. Frontiers in Neuroscience. 13, 95. (2019).

[37] Chaisangmongkon, W., Swaminathan, S. K., Freedman, D. J. & Wang, X.-J. Computing by robust transience: How the fronto-parietal network performs sequential, category-based decisions. Neuron. 93, 1504–1517. (2017).

[38] Denéve, S. & Machens, C. K. Efficient codes and balanced networks. Nature Neuroscience. 19, 375–382. (2016).

[39] Alemi, A., Machens, C. K., Denéve, S. & Slotine, J.-J. E. Learning nonlinear dynamics in efficient, balanced spiking networks using local plasticity rules. In AAAI. (2018).

[40] Zick, J. L., Blackman, R. K., Crowe, D. A., Amirikian, B., DeNicola, A. L., Netoff, T. I. & Chafee, M. V. Blocking NMDAR disrupts spike timing and decouples monkey prefrontal circuits: Implications for activity-dependent disconnection in schizophrenia. Neuron. 98, 1243–1255 (2018).

[41] Shahidi, N., Andrei, A. R., Hu, M. & Dragoi, V. High-order coordination of cortical spiking activity modulates perceptual accuracy. Nature Neuroscience. 22, 1148–1158. (2019).

[42] Ujfalussy, B. B., Makara, J. K., Branco, T. & Lengyel, M. Dendritic nonlinearities are tuned for efficient spike-based computations in cortical circuits. eLife. 4, e10056. (2015).

[43] Yang, G. R., Murray, J. D. & Wang, X.-J. A dendritic disinhibitory circuit mechanism for pathway-specific gating. Nature Communications. 7, 12815 (2016).

